# Taxonomy and phylogeny of the family Suberitidae (Porifera: Demospongiae) in California

**DOI:** 10.1101/2024.01.10.575078

**Authors:** Thomas L. Turner, Greg W. Rouse, Brooke L. Weigel, Carly Janusson, Matthew A. Lemay, Robert W. Thacker

## Abstract

This study presents a comprehensive taxonomic revision of the family Suberitidae (Porifera: Demospongiae) for California, USA. We include the three species previously known from the region, document two additional species previously known from other regions, and formally describe four new species as *Pseudosuberites latke* sp. nov., *Suberites californiana* sp. nov., *Suberites kumeyaay* sp. nov., and *Suberites agaricus* sp. nov. Multi-locus DNA sequence data is presented for seven of the nine species, and was combined with all publicly available data to produce the most comprehensive global phylogeny for the family to date. By integrating morphological and genetic data, we show that morphological characters may be sufficient for regional species identification but are likely inadequate for global classification into genera that reflect the evolutionary history of the family. We therefore propose that DNA sequencing is a critical component to support future taxonomic revisions.

## Introduction

Our understanding of the ecology, evolution, and potential applications of California’s sponge fauna is currently limited by a lack of basic knowledge concerning which species are present, where they occur, how common they are, and how to identify them. Much of what we know about the diversity of California’s sponges comes from seminal taxonomic work conducted 100 years ago and published in a monograph in 1932 (de Laubenfels 1932). Many species are known from only a few individuals, with no information about their abundance and distribution. Later work identified additional species, mainly in the intertidal zone, but rarely published formal descriptions of them. For example, a species of *Suberites* was discovered in the California intertidal by Hartman (1975); despite being a common member of intertidal communities, this species has remained “*Suberites* sp.” in the 50 years since (Bakus & Green 1987; Lee *et al*. 2007). Several species have been discovered and described from the deep ocean (Lee *et al*. 2012; Lundsten *et al*. 2014; Reiswig 2018), but until recently, only a few limited projects have endeavored to sample and identify sponges from the shallow subtidal kelp forest ecosystem (Ristau 1978; Sim & Bakus 1986). Recent publications have revised several orders of California demosponges (Turner 2020, 2021; Turner & Pankey 2023), but many taxa remain unaddressed in the recent literature.

The family Suberitidae contains about 215 valid species across 10 genera (de Voogd, N.J. *et al*. 2023). The evolutionary history of the family has been difficult to discern because very few taxonomically informative morphological characters are known. Combining morphological work with molecular data has proven critical to both suberitid alpha taxonomy (Solé-Cava & Thorpe 1986) and ordinal placement of the family (Morrow & Cárdenas 2015). As shown in the results below, the genera within the family remain highly polyphyletic, so continued integration of genetic and morphological data is likely to be crucial to further progress. The poor preservation of DNA in many (but not all) historical sponge samples has proven to be a roadblock to this integration (Agne *et al*. 2022; Turner & Pankey 2023; Vargas *et al*. 2012). One way forward is via regional revisions that incorporate freshly collected material and combine underwater photography, traditional taxonomic characters, genotyping, and phylogenetic methods. Here, we provide such an analysis for the Suberitidae of California.

Three species of Suberitidae were previously known in California, all of which were known at the time of de Laubenfels’ 1932 monograph, albeit under different generic names in every case: *Suberites latus* Lambe 1893, *Rhizaxinella gadus* (de Laubenfels 1926), and *Protosuberites sisyrnus* (de Laubenfels 1930). North of California, a monograph on the order Hadromerida of British Columbia was published in 2014; at the time of publication, Hadromerida included Suberitidae (Austin *et al*. 2014). This work included *S. latus* and an additional 4 species of Suberitidae that might range into California. To build on these past studies, we have combined existing museum records with fresh collections in Southern and Central California, Oregon, Washington, and British Columbia. In addition to the three species previously known in California, we show that two species known from British Columbia are found in the state, and we formally describe four new species. We combine DNA data from these samples with publicly available genetic data to present an up-to-date global phylogeny of the family.

## Materials & Methods

*Collections.* Sponges were collected from California’s shallow waters and intertidal zone by Thomas Turner. SCUBA-based collection was performed at 77 Southern California locations and 14 locations around the Monterey Peninsula in Central California. The maximum depth investigated was 32 m, but most (75%) sites were less than 22 m deep. Intertidal collections were made at 14 locations from Drake’s Estero (Marin County, Northern California) to Bird Rock (San Diego County, Southern California). Floating docks in 23 harbors and marinas were also investigated, but no Suberitidae were found in these habitats. Collections at each location were focused on novel-looking sponges and sponges that are difficult to identify in the field, while also attempting to photographically document all sponges present. This approach allows presence/absence data to be compiled across sites to form hypotheses about the status and distribution of species investigated, as shown in a supplementary table deposited at Data Dryad (https://doi.org/10.5061/dryad.rbnzs7hhs). Note that the search effort varied across sites, not all species are easily identifiable in the field, and a uniform protocol was not employed; distribution data should therefore be considered non-quantitative and preliminary. Of the 1137 sponge samples collected, 25 were members of the Suberitidae, and include all samples of two of the four species newly described herein.

Samples of *R. gadus* were collected from deep water offshore of Los Angeles by the Schmidt Ocean Institute using the ROV *SuBastian*. A fresh sample of *S. kumeyaay sp. nov.* was collected by Brandon L. Stidum, and archived material from environmental monitoring projects was shared by Megan Lilly and Wendy Enright. Comparative material was collected by the Hakai Institute in British Columbia (8 samples of *S. latus* and one of *S. lambei*); by Jeff Goddard and Gustav Paulay at Cape Arago, Oregon (2 samples of *S. lambei*); by Brooke Weigel on Tatoosh Island, Washington (2 samples of *S. lambei*); and by Thomas Turner in the Olympic Coast National Marine Sanctuary, Washington (4 samples of *S. lambei*).

Previously archived material was acquired on loan from the Royal British Columbia Museum (voucher numbers with RBC), the California Academy of Sciences (voucher numbers with CASIZ), the National History Museum of Los Angeles (voucher numbers with NHMLA), and the Scripps Institution of Oceanography Benthic Invertebrate Collection (vouchers numbers beginning with P). Freshly collected material was deposited in these collections and also the Santa Barbara Museum of Natural History (voucher numbers with SBMNH), the Cheadle Center at the University of California, Santa Barbara (voucher numbers with UCSB), the Hakai Institute (voucher numbers with BHAK), and the Florida Museum (voucher numbers with BULA and UF). Voucher numbers are listed in the systematics section, and are available in tabular format in the supplementary materials deposited at Data Dryad (https://doi.org/10.5061/dryad.rbnzs7hhs).

### Morphology

Spicules were examined after digesting sponge subsamples in bleach. Scanning electron images were taken with a FEI Quanta400F Mk2 after coating spicules with 20 nm of carbon. Skeletal architecture was interrogated by hand cutting tissue sections and digesting them in one of two ways. Some were moved from 95% ethanol to a 100% ethanol bath, then cleared using Histoclear (National Diagnostics). Others were digested in a mixture of 97% Nuclei Lysis Solution (Promega; from the Wizard Genomic DNA Purification kit) and 3% 20mg/ml Proteinase K (Promega). This digestion eliminates cellular material while leaving the spongin network intact. Spicule measurements were made on images using ImageJ (Schneider et al. 2012) after calculating the number of pixels per mm with a calibration slide. Spicule length was measured as the longest possible straight line from tip to tip, even when spicules were curved or bent. Spicule width was measured at the widest point, excluding adornments like swollen tyles. Scale bars on figures are precise for microscope images but often approximate for field photos. All spicule measurements are available as raw data at Data Dryad (https://doi.org/10.5061/dryad.rbnzs7hhs).

### Genotyping

DNA was extracted with several different kits including the Wizard Genomic DNA Purification kit (Promega) and the Qiagen Blood & Tissue kit. Three primer sets were used to sequence a fragment of the *cox1* mitochondrial locus. Most samples were genotyped at a ∼1200 bp fragment using the following primers (LCO1490: 5-GGT CAA CAA ATC ATA AAG AYA TYG G-3’; COx1-R1: 5’-TGT TGR GGG AAA AAR GTT AAA TT-3’); these amplify the “Folmer” barcoding region and the “co1-ext” region used by some sponge barcoding projects (Rot *et al*. 2006). When amplification with these primers failed, the Folmer region alone was amplified from some samples using the following primers: (LCO1490: 5’-GGT CAA CAA ATC ATA AAG AYA TYG G-3’; HCO2198: 5’-TAA ACT TCA GGG TGA CCA AAR AAY CA-3’) (Folmer *et al*. 1994). Samples sequenced at the Hakai Institute were instead amplified with the primers jgLCO1490: 5’-TIT CIA CIA AYC AYA ARG AYA TTG G-’3; jgHCO2198: 5’-TAI ACY TCI GGR TGI CCR AAR AAY CA-3’ (Geller *et al*. 2013). The D1–D5 region of the *28S* rDNA nuclear locus was sequenced by combining two amplicons: the ∼800 bp D1-D2 region using primers Por28S-15F (5’-GCG AGA TCA CCY GCT GAA T-3’) and Por28S-878R (5’-CAC TCC TTG GTC CGT GTT TC-3’) and the ∼800 bp D3-D5 region using primers Por28S-830F (5’-CAT CCG ACC CGT CTT GAA-3’) and Por28S-1520R (5’-GCT AGT TGA TTC GGC AGG TG-3’) (Morrow *et al*. 2012). Samples sequenced at the Hakai Institute were instead amplified at overlapping 28S regions by combining the following primers, 28F63mod: 5’-ACC CGC TGA AYT TAA GCA TAT HAN TMA G-3’; 28R077sq: 5’-GAG CCA ATC CTT WTC CCG ARG TT-3’ and 28F1411: 5’-CCG CTA AGG AGT GTG TAA CAA C-3’; 28RampRC: 5’-ACC TGT CTC ACG ACG KTC TRA ACC CAG CTC-3’ using the protocols described by Thacker *et al*. (2013).

PCR was performed using the following conditions. For the *cox1* locus with the Rot primers: 95°C for 3 min, followed by 35 cycles of 94°C for 30 sec, 50°C for 30 sec, 72°C for 75 seconds, followed by 72°C for 5 minutes; for the *cox1* locus with Folmer primers, the annealing temperature was modified to 52°C with 60 second extension time; for both *28S* amplicons, a 53°C annealing temperature and 60 second extension time was used. PCR was performed in 50 μl reactions using the following recipe: 24 μl nuclease-free water, 10 μl 5x PCR buffer (Gotaq flexi, Promega), 8 μl MgCl, 1 μl 10mM dNTPs (Promega), 2.5 μl of each primer at 10 μM, 0.75 bovine serum albumin (10 mg/ml, final conc 0.15 mg/ml), 0.25 μl Taq (Gotaq flexi, Promega), 1 μl template. ExoSAP-IT (Applied Biosystems) was used to clean PCRs, which were then sequenced by Functional Biosciences using Big Dye V3.1 on ABI 3730xl instruments. Blastn was used to verify that the resulting traces were of sponge origin. All sequences have been deposited in GenBank; accession numbers are shown in the phylogenies and also listed in the supplementary table in Data Dryad (https://doi.org/10.5061/dryad.rbnzs7hhs).

### Phylogenetic methods

We used the NCBI taxonomy browser and blast to assemble all available suberitid sequences and comparative sequences from the Halichondriidae and outgroups. These data include unpublished datasets and numerous previous studies (Abe *et al*. 2012; Becking *et al*. 2013; Ereskovsky *et al*. 2018; Erpenbeck *et al*. 2007, 2012; Idan *et al*. 2018; Lukić-Bilela *et al*. 2008; Melis *et al*. 2016; Morrow *et al*. 2012, 2013, 2019; Najafi *et al*. 2018; Nichols 2005; Núñez Pons *et al*. 2017; Pankey *et al*. 2022; Perovic-Ottstadt *et al*. 2004; Samaai *et al*. 2017; Thacker *et al*. 2013; Turner & Lonhart 2023; Vargas *et al*. 2015; Vicente *et al*. 2022). For *cox1*, sequences were excluded if they did not include the Folmer barcoding region. For *28S*, sequences were excluded if they did not include the C2-D2 barcoding region. Sequences were aligned using MAFFT v.7 online (Katoh *et al*. 2017), and alignment files are available as supplementary data from Data Dryad. Phylogenies were estimated with maximum likelihood in IQ-Tree (Nguyen *et al*. 2015; Trifinopoulos *et al*. 2016); the ultrafast bootstrap and Shimodaira– Hasegawa approximate likelihood ratio test (SH-aLRT) were used to measure node confidence (Hoang *et al*. 2018). ModelFinder (Kalyaanamoorthy *et al*. 2017) was used to choose the optimum model for each tree based on Bayesian Information Criterion; TIM3+F+R3 model for both loci. Figures were produced by exporting IQ-Tree files to the Interactive Tree of Life webserver (Letunic & Bork 2019).

## Results

### Molecular phylogenies

We constructed phylogenies of the *28S* nuclear locus (figure 1) and the *cox1* mitochondrial locus (figure 2) separately in order to include as many taxa as possible. These are the most frequently genotyped loci in Porifera, and these phylogenies are the most comprehensive to date for the order Suberitida. A recently published super matrix analysis — with more genetic data but many fewer species — is highly congruent with our phylogenies (Pankey *et al*. 2022).

**Figure 1.**
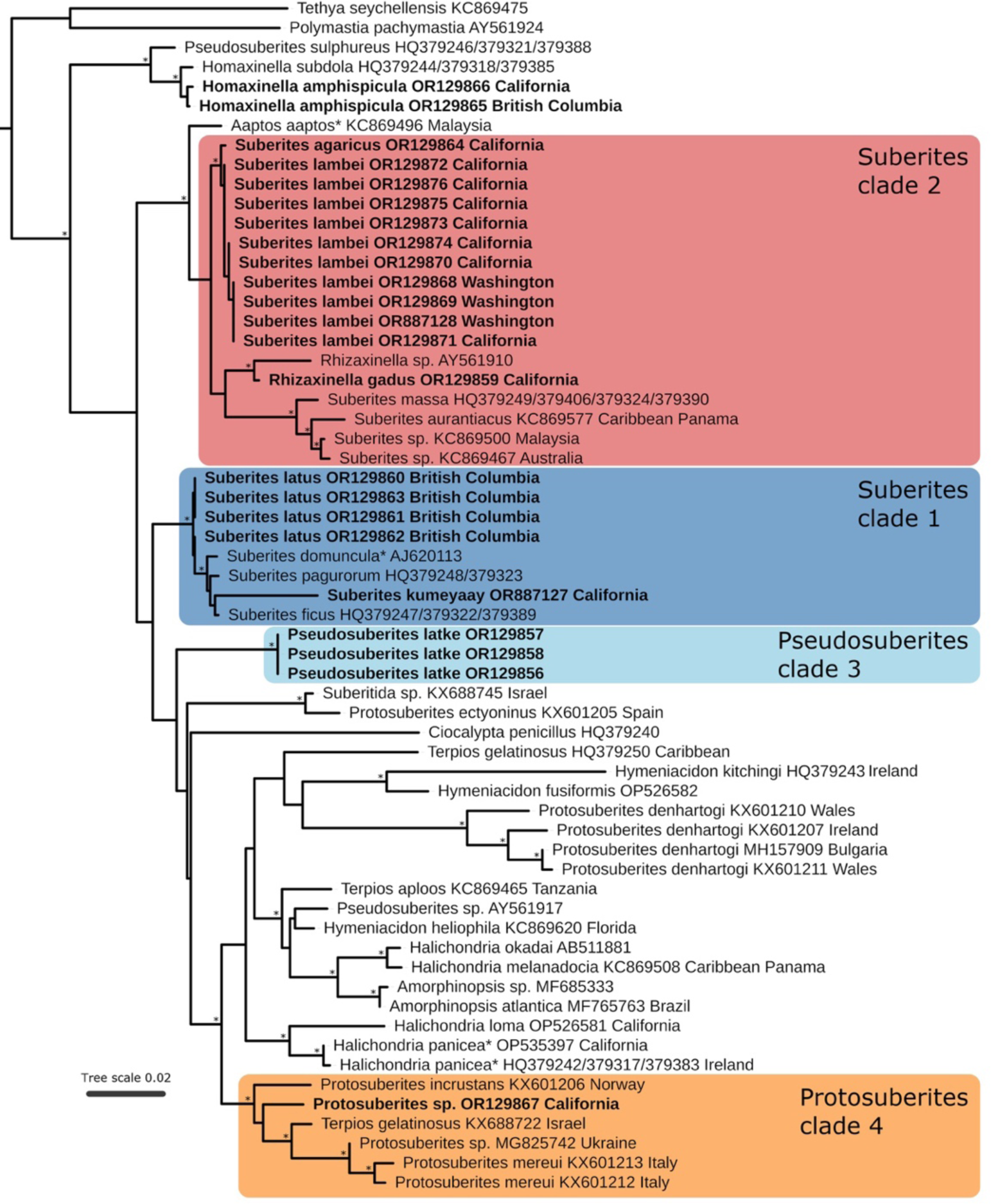
Maximum likelihood phylogeny of the *28S* locus. Genbank accession numbers and collection locations (when known) are shown; bold indicates new sequences. Asterisks at nodes indicate SH-aLRT ≥ 80% and UFbootstrap ≥ 95%. Scale bar indicates substitutions per site. Asterisks in node names indicate type species of a genus.

**Figure 2.**
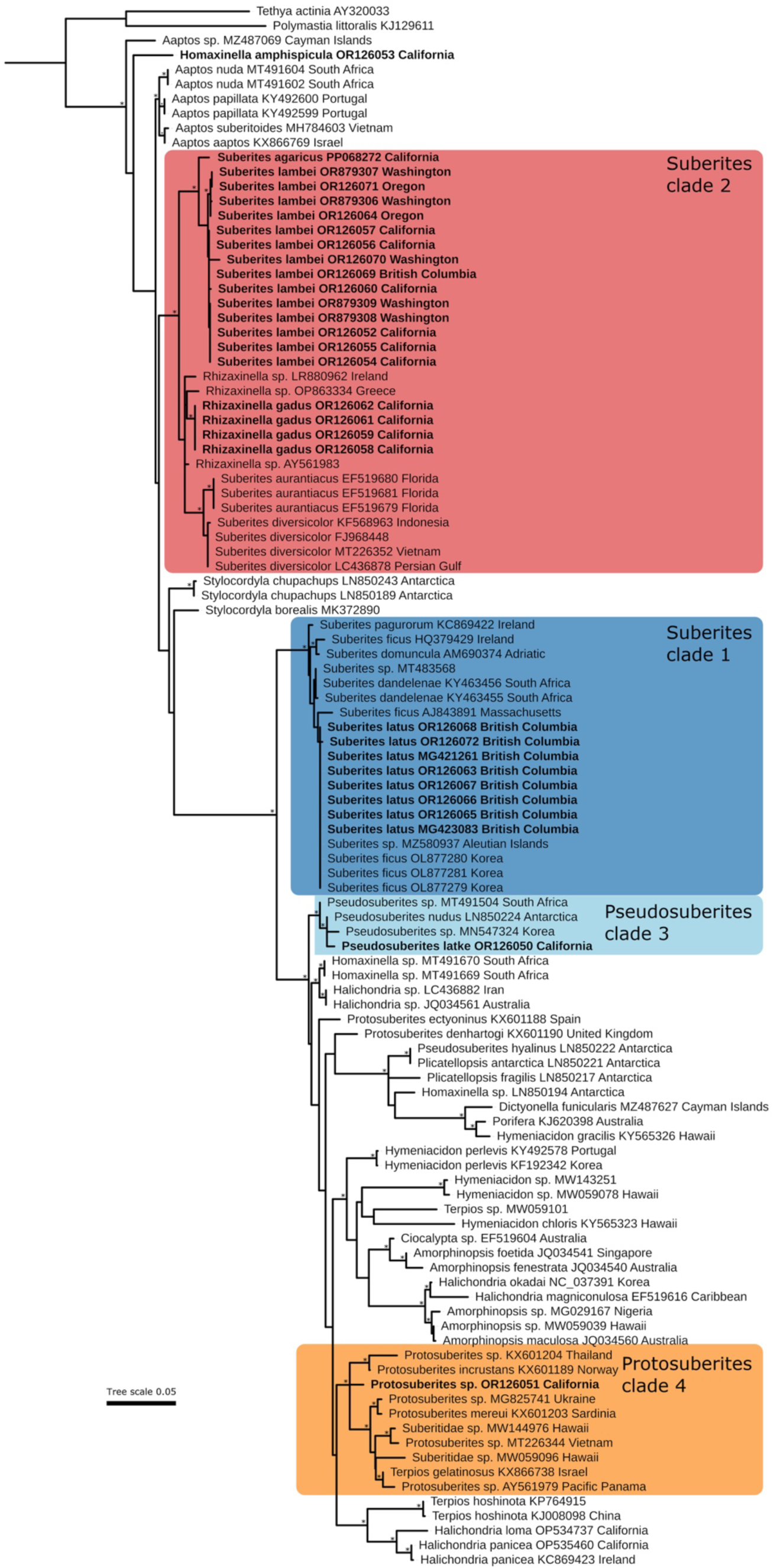
Maximum likelihood phylogeny of the *cox1* locus. Genbank accession numbers and collection locations (when known) are shown; bold indicates new sequences. Asterisks at nodes indicate SH-aLRT ≥ 80% and UFbootstrap ≥ 95%. Scale bar indicates substitutions per site. Asterisks in node names indicate type species of a genus.

The type species *Suberites domuncula* (Olivi 1792) forms a highly supported clade containing a world-wide sample of other *Suberites* (clade 1, blue in figures 1 & 2). This clade includes the species known to be associated with hermit crabs, including *Suberites latus*, the hermit crab-associated representative from the North Pacific. There is a deep divergence between this *Suberites ‘sensu stricto’* clade and clade 2, shown in red, which contains the remaining *Suberites* and the genus *Rhizaxinella*. *Rhizaxinella* are differentiated from *Suberites* by their stalked morphology: data from these two loci are insufficient to resolve whether stalks evolved only once in this clade. It is clear, however, that the genus *Suberites* contains at least two well-differentiated clades, and it seems unlikely that spicule-based morphological characters will be able to resolve membership in these clades.

The North Pacific *Homaxinella amphispicula* (de Laubenfels 1961) is very morphologically similar to the North Atlantic *H. subdola* (Bowerbank 1866), and DNA supports a close relationship at the 28S locus. The placement of the genera *Homaxinella* and *Aaptos* are inconsistent between loci, and will require additional data to resolve.

The remaining California suberitid species appear more closely related to the Halichondriidae than to *Suberites*. *Pseudosuberites latke* sp. nov. forms a clade with *Pseudosuberites* from South Africa, Antarctica, and Korea at the *cox1* locus (clade 3, light blue). The type species of the genus, *P. hyalinus* Ridley & Dendy 1887, is found in a different location in the *cox1* tree, but this result should be considered tentative. The sequence is identical to a sample from the same collection that is identified as a *Plicatellopsis*, which may indicate some cross-contamination of DNA in that study. Several other samples identified as *Pseudosuberites* are found in our *28S* tree, however, and also do not form a clade with the new species. This genus is therefore likely polyphyletic and in need of revision.

Thinly encrusting Suberitidae are placed in either *Terpios* (which have a gelatinous consistency, lobate tylostyles & low spicule density) or *Protosuberites* (with upright tylostyles) (Morrow & Cárdenas 2015; van Soest 2002). The majority of *Protosuberites* fall within a single clade, shown in orange (clade 4, orange). The type species for the genus has not been sequenced, however, and the genus is polyphyletic due to *Protosuberites denhartogi* (van Soest & De Kluijver 2003) and *Protosuberites ectyoninus* (Topsent 1900) falling in other parts of the tree. The difficulty of identification in these morphologically simple sponges is highlighted by the observations that *P. denhartogi* appears to be a species complex based on *28S* data, and samples identified as *Terpios gelatinosa* (Bowerbank 1866) fall in two different clades.

### Systematics

#### Summary of California Suberitidae

Genus Suberites

*Suberites latus* Lambe 1893
*Suberites californiana* sp. nov. Turner 2023
*Suberites kumeyaay* sp. nov. Turner 2023
*Suberites lambei* Austin, Ott, Reiswig, Romagosa & McDaniel, 2014
*Suberites agaricus* sp. nov. Turner 2023

Genus Rhizaxinella

*Rhizaxinella gadus* (de Laubenfels 1926)

Genus Pseudosuberites

*Pseudosuberites latke* sp. nov. Turner 2023

Genus Protosuberites

*Protosuberites sisyrnus* (de Laubenfels 1930)
*Protosuberites sp*.

Genus Homaxinella

*Homaxinella amphispicula* (de Laubenfels 1961)

**Genus Suberites Nardo, 1833**

***Suberites latus* Lambe, 1893**

Figures 1-3

**Figure 3.**
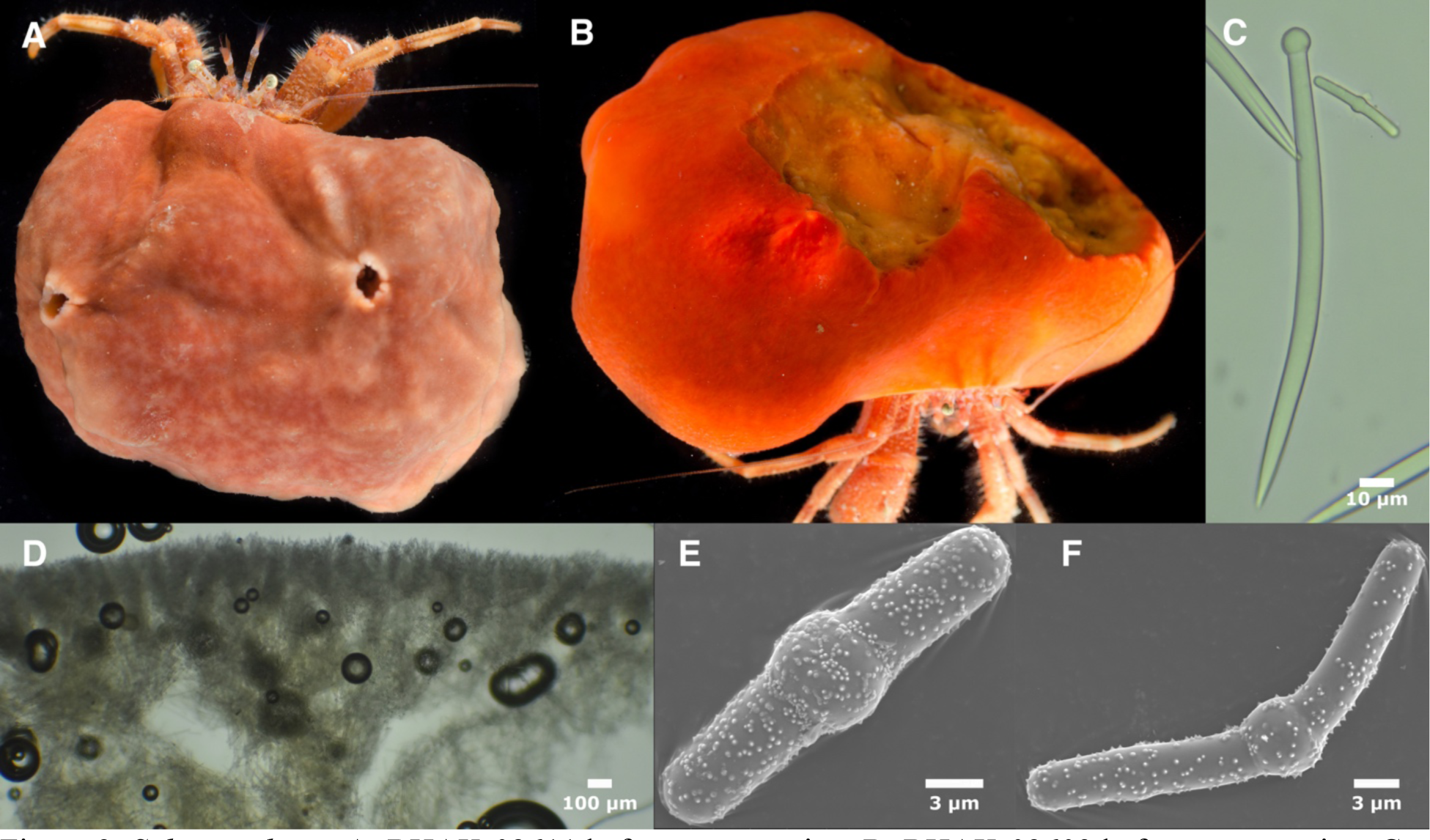
*Suberites latus*. A: BHAK-02611 before preservation. B: BHAK-02603 before preservation. C: Tylostyle and centrotylote microstrongyle. D: Cross-section showing ecotosomal brushes and confused inteorior. E-F: centrotylote microstrongyles. C-F from Rattenbury Pinnacle sample.

### Synonyms

*Ficulina suberea var. lata* (de Laubenfels 1932)
*Choanites suberea var. lata* (de Laubenfels 1961)
*Suberites ficus* (Bakus & Green 1987)

#### Material examined

UF4128/BHAK-02603, UF4127/BHAK-02601, UF47673/BHAK-02604, UF4129/BHAK-02605 & UF4131/BHAK-02611, all collected at Rattenbury Pinnacle, British Columbia, Canada, (51.691, −120.080), 21 m, 8/3/17; UF4143/BHAK-03126, Big Spring Creek, British Columbia, Canada, (51.6501, −128.069), 2 m, 8/6/17; BHAK-10221 & BHAK-10390, both collected at Wedgeborough Point, British Columbia, Canada, (51.647, −127.956), 15 m, 5/27/19.

#### Morphology

Massive, amorphous sponges, nearly always containing a hermit crab. Color in life variable from vivid orange to reddish brown. Firm and smooth to the touch.

#### Skeleton

Ectosomal skeleton of tylostyles in upright bouquets. Tylostyles in choanosome in confusion with no apparent pattern.

#### Spicules

Tylostyles in two size classes, with spined centrotylote strongyles as microscleres. Short tylostyles are restricted to ectosome, where some long tylostyles can also be found. Tylostyle size classes divided at 200 μm following Austin et al. (2014). Data are from two individuals from the Rattenbury Pinnacle collection, which were loaned in a common container.

Long tylostyles: those we examined were nearly always smoothly curved or with a slight bend near the center (figure 3C), but a previous report found them to sometimes be straight or sinuous (Austin et al. 2014). Most taper abruptly to a sharp point, but rounded points or strongylote endings present on a substantial minority. Heads usually terminal, round, and symmetrical, but occasionally subterminal. Shafts are often slightly fusiform, with thickest section in center (figure 3C), but some have consistent thickness throughout. 216–305–409 x 3–7–11 μm (n=95)

Short tylostyles: shaped the same as long tylostyles. 103–158–198 x 2–7–9 μm (n=67) All tylostyles considered together: 103–245–409 x 2–7–11 μm (n=162), modes at approximately 170 and 310 μm.

Centrotylote microstrongyles: variable abundance from rare to common, and apparently sometimes absent. Covered in very small spines which are difficult to see in a light microscope (figure 3C). Tylote ball is generally near the center but sometimes at one end. Shaft sometimes straight but often curved or bent at the center (figure 3F). Austin et al. (2014) report that these spicules are choanosomal. Consistent with this report, they were much more common in the choanosome in one sample we examined, and very rare in both choanosome and ectosome in the other sample we examined. 14–24–33 x 1–4–6 μm (n=84 length, n=34 width).

#### Distribution and habitat

This species is known from a wide depth range (6–183 m) from the Aleutian Islands, Alaska to British Columbia (Austin *et al*. 2014). It is nearly always found in an association with a hermit crab, either on a gastropod shell or in the shape of a gastropod shell that has presumably dissolved post-settlement. Genetic data from Korean samples suggest the range extends from the Aleutians into the Northwest Pacific as well (see remarks). The presence of this species in California is unconfirmed; if present, it is likely only present in deep water (>100 m; see remarks).

#### Remarks

This species was described from samples collected near Vancouver Island (Pacific Canada), but the morphological similarity to other *Suberites* has led to uncertainty regarding its species status (Austin *et al*. 2014). A recent revision acknowledged the difficulty of morphological identification in this group, but supported the concept of *S. latus* as a North Pacific relative of the North Atlantic *S. ficus* complex, and went on to propose that “this species complex would be a good candidate for DNA barcoding” (Austin *et al*. 2014). Here we present DNA data from samples collected in British Columbia, near the type location. These samples are genetically differentiated from morphologically similar Atlantic species such as *S. ficus* Johnson, 1842 and *S. pagurorum* Solé-Cava & Thorpe, 1986, supporting species status for *S. latus*. It is also notable that the microstrongyles of this species are more common in the choanosome, which contrasts with other *Suberites*, where they are more common in the ectosome.

We were unable to confirm the presence of *S. latus* in California. All freshly collected and previously vouchered material from California’s shallow waters was found to be either *S. kumeyaay* sp. nov. or *S. lambei*. Consistent with the lack of vouchered samples from shallow water in California, amateur observations of *Suberites* with hermit crabs on the site iNaturalist extend from Kodiak, Alaska to the San Juan Islands in northern Washington, but are absent farther South. *Suberites* associated with hermit crabs are found in deep water in Southern California, by environmental monitoring projects, nearly always below 100 m (Megan Lilly, City of San Diego, *pers. comm.*). Well-characterized samples are morphologically differentiated from the Northern *S. latus*, and we allocate them to *Suberites californiana* sp. nov., described below. However, it remains possible that *S. latus* also ranges into California in deep water, as few deep-water California samples are available for review.

At the *cox1* locus, Genbank records from Korea, previously identified as *S. ficus*, are genetically identical to *S. latus*, raising the possibility that the species range extends from the Aleutians into the Northwest Pacific (figure 2). Confirmation would require DNA sequence from additional loci and morphological data, as the slow rate of evolution at *cox1* in sponges sometimes leads to identical sequences between populations that are good species under most species concepts (Huang *et al*. 2008; López-Legentil *et al*. 2010; Pöppe *et al*. 2010; Turner & Pankey 2023). Shallow samples from Peru were recently described as *Suberites cf. latus* (Cóndor-Luján *et al*. 2023). Due to the disjunct distribution, and lack of association with hermit crabs, we think it likely that DNA data would reveal these to be a different species.

***Suberites californiana* sp. nov. Turner 2023**

Figure 4

**Figure 4.**
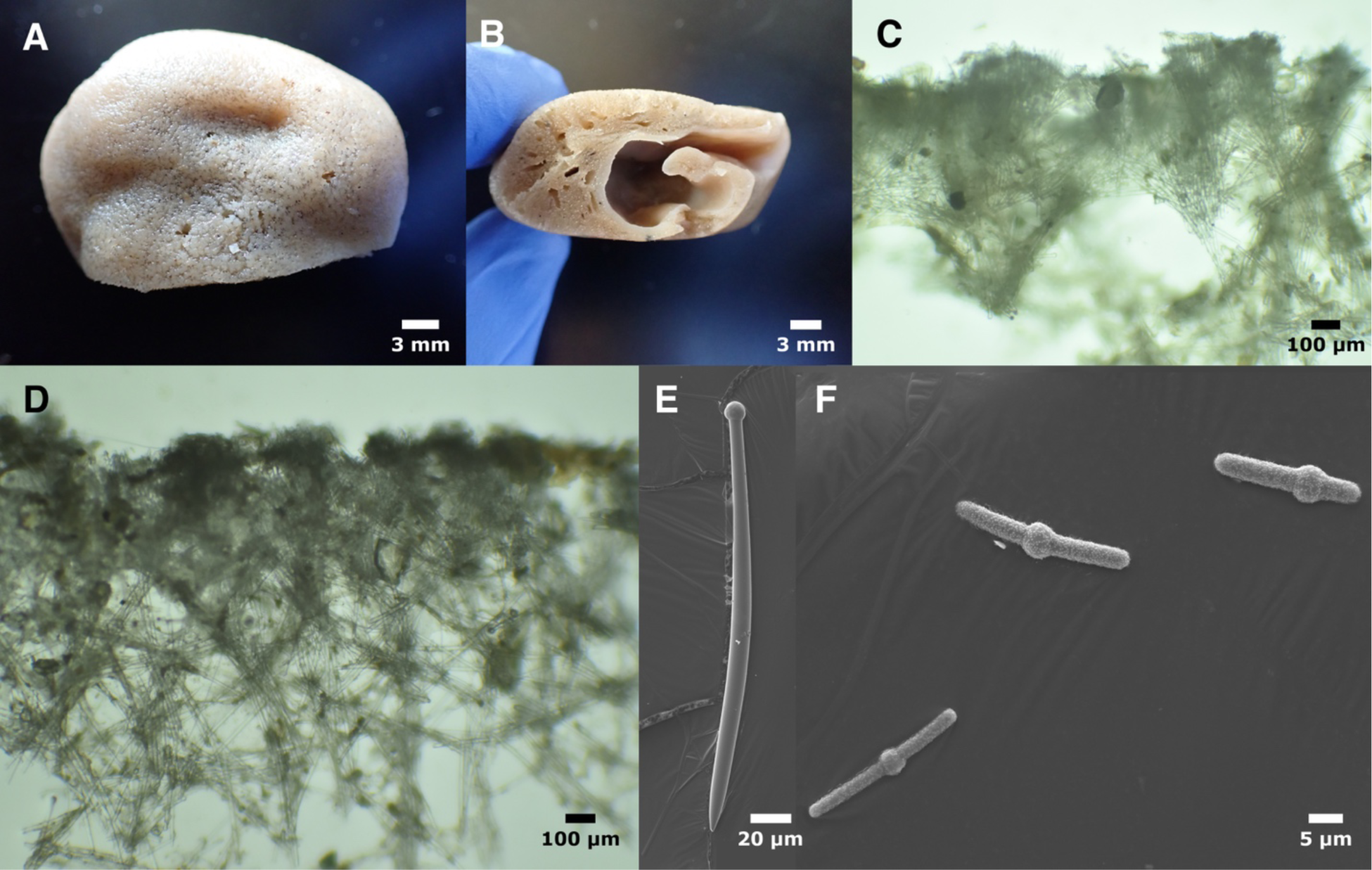
*Suberites californiana.* Holotype, post-preservation, from above (A) and the side (B), the latter sectioned to show hermit crab cavity. C: Cross-section showing ectosomal brushes. D: Cross-section showing ecotosomal brushes and confused interior. E: Tylostyle. F: Centotylote microsontrongyles. All photos of holotype.

#### Material examined

Holotype: P362, Near Anacapa Island, (34.00, −119.39), 223-242 m, 4/3/52.

#### Etymology

Named for the state of California.

#### Morphology

Massive, roughly globular, 3 cm across, with a spiral cavity for a hermit crab (figure 4B). Color in life not recorded; beige preserved. Firm and smooth to the touch. A previously described sample was growing on a gastropod shell, but it was not stated whether the shell contained a hermit crab (Bakus & Green 1987).

#### Skeleton

Choanosomal skeleton is a confused lattice of single tylostyles and tylostyle bundles (figure 4D). Spiculation becomes denser approaching the ectosome, and while still confused, upright bouquets typical of the genus are also apparent (figure 4C). A few microstrongyles are found scattered in the choanosome, but are much more abundant at sponge surface.

#### Spicules

Tylostyles in two size classes, with spined centrotylote strongyles as microscleres. Ectosomal tylostyles are nearly all in the short size class, while choanosomal tylostyles are nearly all in the large size class.

Long tylostyles: usually slightly curved or bent near the center, but occasionally straight or sinuous (figure 4E). Most taper abruptly to a sharp point, but rounded points or strongylote endings present. Heads terminal, round, and symmetrical, but occasionally subterminal. Many are slightly fusiform, with thickest section in center (figure 4E), but others have consistent thickness throughout. 408–574–693 x 10–14–20 μm (n=46).

Short tylostyles: shaped the same as long tylostyles. 155–250–395 x 8–11–16 μm (n=41).

All tylostyles considered together: 155–421–693 x 8–13–20 μm (n=87), modes at approximately 250 and 600 μm.

Centrotylote microstrongyles: microspined, but appearing smooth under light microscopy. Generally tylote in or near the center (figure 4F), but sometimes at one end. Spicules slightly curved or straight. 13–23–52 x 2–3–5 μm (n=55; measurements includes a single much longer spicule, perhaps an aberration, but nearly all are between 13 and 32 μm in length).

#### Distribution and habitat

Only two samples can be confidently assigned to this species, both from deep water (183 m – 242 m) in Southern California.

#### Remarks

*Suberites latus* from the Pacific Northwest (Alaska, British Columbia, Washington) have spicules with fairly consistent dimensions. Combined, three previous analyses have found maximum tylostyle lengths between 307 and 409 μm. The two samples analyzed here had maximum tylostyle lengths of 354 and 409 μm; seven additional samples, collected at depths varying from <20 m to 126 m, had maximum lengths between 307 and 394 μm (Austin *et al*. 2014); and 60 individuals from Alaska had maximum lengths to 406 μm (Lambe 1895). In contrast, both samples known from Southern California have much longer spicules. The sample analyzed here had long tylostyles with a maximum length of 693 μm (∼70% longer than *S. latus*). Quantitative data has been published for one other sample: a *Suberites* (previously identified as *S. ficus*) growing on a gastropod shell at 183 m (Bakus & Green 1987). This sample had spicules with very similar dimensions as the sample analyzed here, with a maximum length of 680 μm. In addition, the sample analyzed here had microscleres that were abundant in the ectosome, in contrast to the rarity and/or choanosomal distribution in *S. latus* (the location of the microscleres was not specified in the previously analyzed sample). We therefore designate all *Suberites* in California with microscleres and tylostyles longer than 500 μm to be *Suberites californiana* sp. nov. Additional morphological and genetic work on deep *Suberites* is needed to determine if *S. latus* is also found in the state.

***Suberites kumeyaay* sp. nov. Turner 2023**

Figures 1, 5

**Figure 5.**
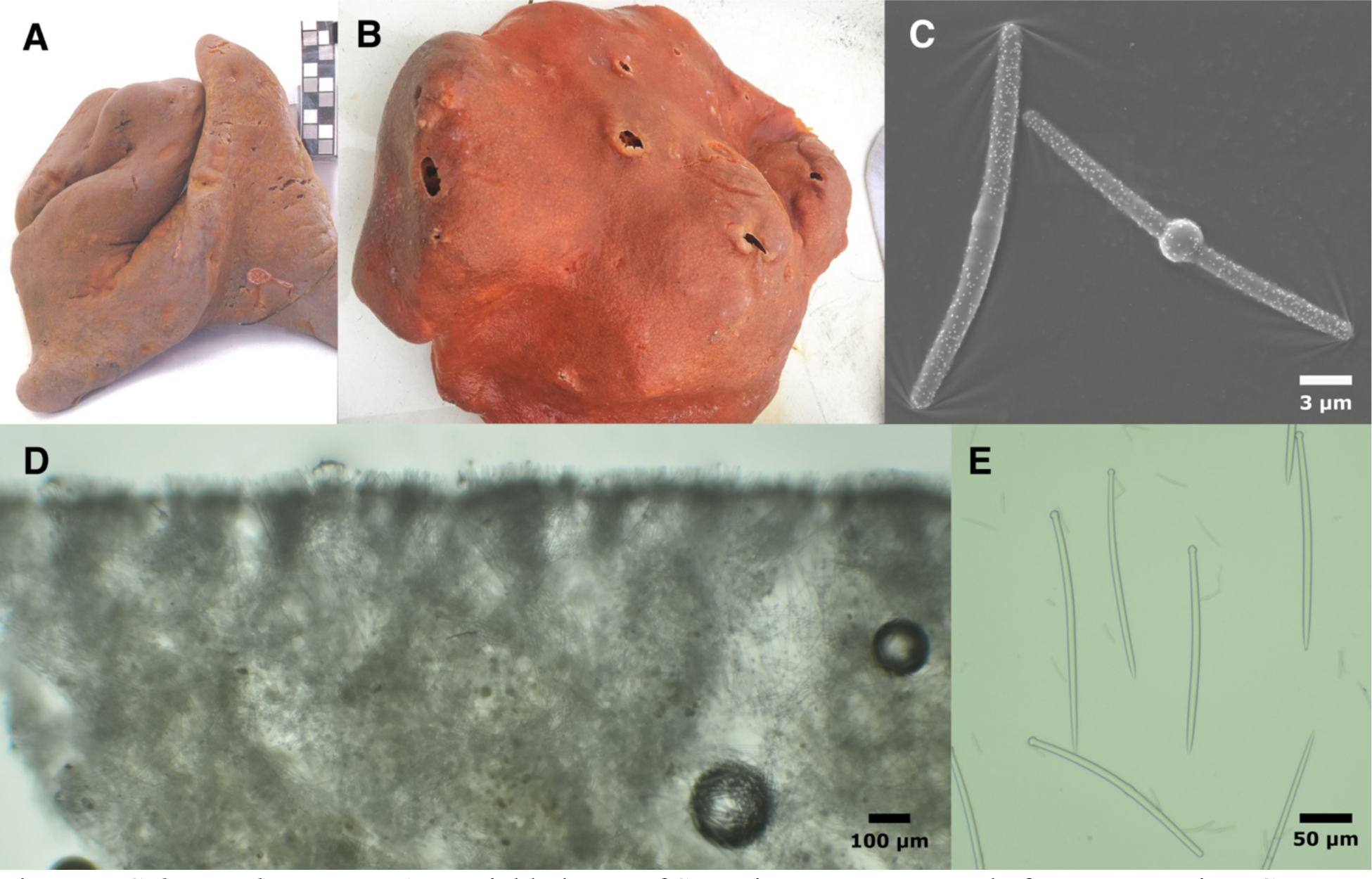
*Suberites kumeyaay*. A-B: Field photos of San Diego Bay sponges before preservation. C: centrotylote microstrongyles. D: Cross-section showing ecotosomal brushes and confused interior. E: example tylostyles, including one with slightly off-center head (left), slightly non-terminal head (center) and rare centrotylote strongyle modification (bottom). C-E from holotype.

#### Material examined

Holotype: TLT1439, San Diego Bay, 14.25 m, 8/31/23; Paratypes: TLT1340, Offshore San Diego, (32.53197, −117.18800), 31 m, 7/25/17; TLT1339, San Diego Bay, 10 m, 8/17/98; TLT1341, San Diego Bay, 10 m, 8/13/98; TLT1441, San Diego Bay, 10 m, 8/14/98; TLT1442, San Diego Bay, 10 m, 8/24/98.

#### Etymology

Named for the Kumeyaay people, who have inhabited the San Diego area since pre-Columbian times.

#### Morphology

Massive, amorphous, lobate sponges. Bright orange to brownish orange alive, light brown preserved. The holotype is a roughly triangular lobe cut from a larger sponge, 9 x 7 x 2 cm across. It has scattered oscula, 1-5 mm in diameter. Dried samples available for examination measure up to 22 x 10 cm, but field photos indicate larger sponges are sometimes found. The holotype was preserved in 95% ethanol, but other samples were preserved in formalin and/or dried.

#### Skeleton

Ectosomal skeleton of tylostyles in upright bouquets (figure 5D), with microstrongyles among the bouquets. Choanosomal skeleton of tylostyles in confusion with no apparent pattern. A few microstrongyles are found scattered in the choanosome, but are much less abundant than at sponge surface.

#### Spicules

Long and short tylostyles, with spined centrotylote strongyles as microscleres. Tylostyles: usually slightly curved and weakly fusiform (thickest in center), but longest tylostyles are often straight. Heads are typically round and terminal, but frequently off-center and sometimes sub-terminal (figure 5E). The length distribution is bimodal, but with many styles of intermediate size that make it difficult to categorize spicules as short vs. long. Choanosomal styles average 60% longer than ectosomal styles.

Holotype. Ectosome, 121–193–272 x 5–7–10 μm (n=40); choanosome, 151–260–305 x 1–7–9 μm (n=40); both domains combined, 121–226–305 x 1–7–10 μm (n=80).

All samples pooled, 81–223–324 x 1–7–13 μm (n=338), with modes at 135 and 285 μm.

Centrotylote microstrongyles: microspined, but appearing smooth under light microscopy. Swollen center variable from well-developed to barely present (figure 5C); it is generally in the center but sometimes off-center or at one end. Spicules usually slightly curved but sometimes straight. Holotype: 13–29–45 x 1–2–3 μm (n=97 length, n=35 width). All samples pooled: 13– 28–45 x 1–2–4 μm (n=175 length, n=112 width).

#### Distribution and habitat

Common in the shallow waters of San Diego Bay in Southern California. One sample was also found 6 km offshore of San Diego in 31 m of water.

#### Remarks

Environmental monitoring projects in San Diego Bay, in extreme southern California, frequently yield large *Suberites* that are not associated with gastropod shells or hermit crabs; these were previously referred to as “Porifera sp. SD 4” by the Southern California Association of Marine Invertebrate Taxonomists (Megan Lilly, City of San Diego, *pers. comm.*). These differ from *Suberites californiana* sp. nov. in that they are 1) not associated with gastropod shells and hermit crabs, 2) found in shallow water (4–31 m, vs. 180–242 m for *S. californiana*), 3) have much shorter tylostyles (modes at 135 and 285 μm vs. 250 and 600 μm for *S. californiana*; maximum lengths 305–324 μm vs. 600–693 μm for *S. californiana*), and 4) possess microstrongyles that are slightly, but significantly (p<0.001) longer than *S. californiana* (mean = 28 μm vs. 23 μm for *S. californiana*). *Suberites kumeyaay* sp. nov. is also morphologically distinguished from *S. latus* in that they are 1) not associated with gastropod shells and hermit crabs, 2) have abundant ectosomal microscleres (vs. rare and/or choanosomal in *S. latus*), 3) have slightly shorter tylostyles (modes at 135 and 285 μm vs. 170 and 310 μm for *S. latus*; maximum lengths 305–324 μm vs. 354–409 μm for *S. latus*), and 4) possess microstrongyles that are slightly, but significantly (p<0.001) longer than *S. latus* (mean = 28 μm vs. 24 μm for *S. latus*). It is easily differentiated from all other *Suberites* in the region, which lack microscleres.

Previously collected samples of *S. kumeyaay* sp. nov. were either preserved in formalin or dried, but we were able to generate 28S sequence from a freshly collected sample, preserved in ethanol, graciously collected for this purpose by Brandon L. Stidum. This 28S sequence is most similar to *S. ficus*, a North Atlantic species that is very morphologically similar to *S. kumeyaay* sp. nov. These species are quite genetically divergent, however, with 3.60% pairwise sequence differences, which is much higher than the sympatric, reproductively isolated *S. ficus* and *S. pagorum* (0.84% divergence). Combined with their highly disjunct distribution, this leads to confidence in assigning Southern California samples a new species.

***Suberites lambei* Austin, Ott, Reiswig, Romagosa & McDaniel 2014**

Figures 1-2, 6

**Figure 6.**
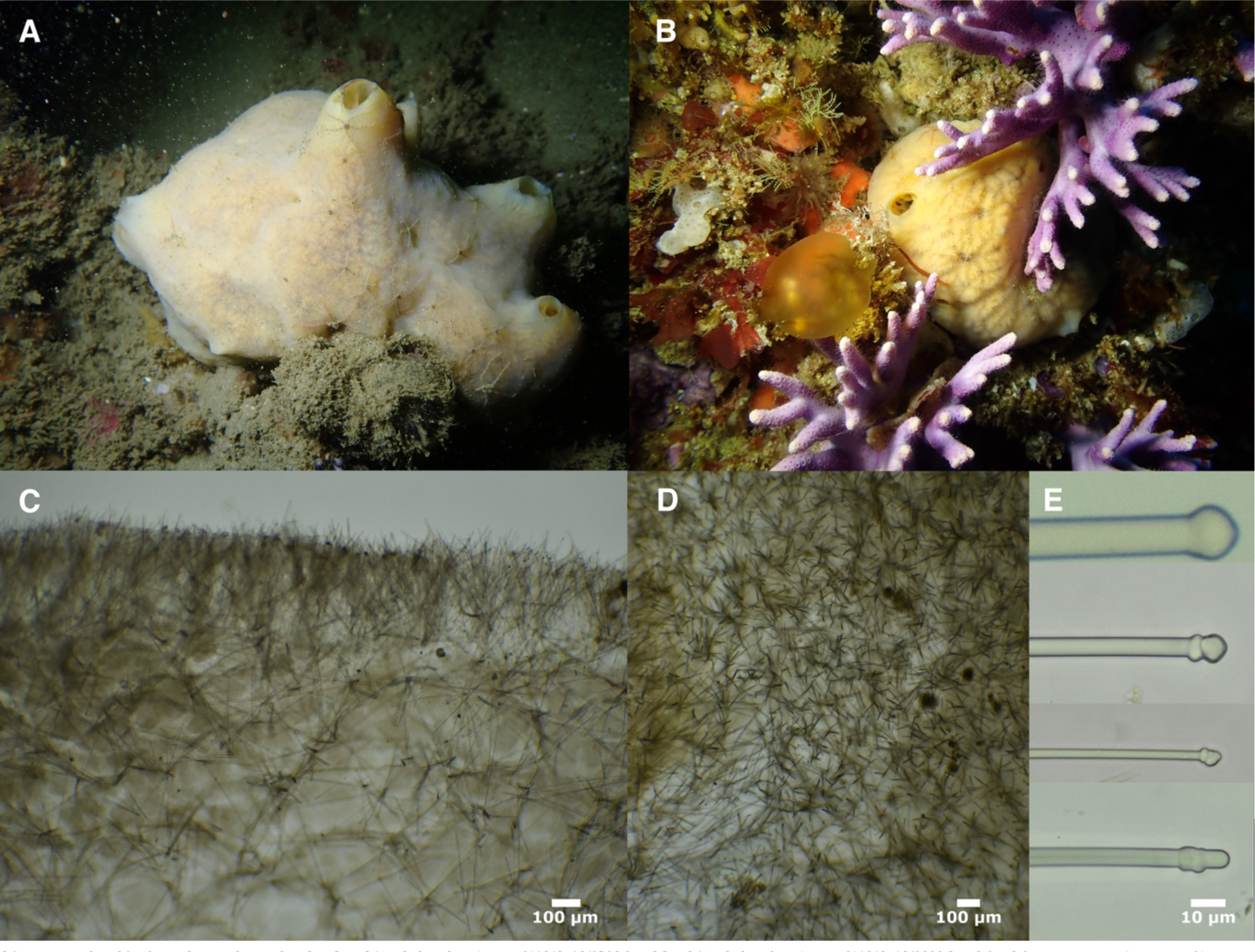
*Suberites lambei.* A: Field photo of TLT296. B: Field photo of TLT991. C: Cross-section of TLT432 showing ectosomal brushes and confused choanosomal skeleton. D: Tangential section of TLT432 showing ecotosomal skeleton at surface. E: Variation in spicule width and head shape; thick tylostyle with well-formed head from TLT684, thin spicules with lumpy heads from TLT1270.

### Synonyms

Suberites sp. (Bakus & Green 1987; Hartman 1975; Lee *et al*. 2007)

#### Material examined

Holotype: RBC982-00066-001, Barkley Sound, George Frasier Island, British Columbia, (48.90833,-125.51333), intertidal, 7/29/76; Paratype: RBC982-00069-001, Gowlland Pt., S. Pender Island, British Columbia, (48.73667,-123.18167), 3 m, 3/18/77; CASIZ302850, South Shore, San Nicolas Island, San Nicolas Is., (33.21717,-119.47450), 10 m, 9/28/11; CASIZ302796, The Arch, San Clemente Island, San Clemente Is., (32.96000,-118.51533), 7 m, 9/19/11; TLT192, Arroyo Hondo, Santa Barbara, (34.47182,-120.14262), 4-8 m, 7/29/19; TLT441, Elwood Reef, Santa Barbara, (34.41775,-119.90150), 9-15 m, 10/23/19; TLT374, Wreck of the Cuba, San Miguel Is., (34.03009,-120.45575), 6-12 m, 10/6/19; BULA0547/NHMLA16557, Halfway Reef, Los Angeles, (33.76265,-118.42560), 15-23 m, 8/23/19; BULA0549/NHMLA16559, Halfway Reef, Los Angeles, (33.76265,-118.42560), 15-23 m, 8/23/19; TLT432, Elwood Reef, Santa Barbara, (34.41775,-119.90150), 9-15 m, 10/23/19; TLT672, La Jolla Cove, San Diego, (32.85227,-117.27239), 15 m, 8/14/20; TLT684, Hazard Canyon, San Luis Obispo, (35.28959,-120.88415), intertidal, 12/12/20; TLT660, Hazard Canyon, San Luis Obispo, (35.28959,-120.88415), intertidal, 12/12/20; TLT825, Cave Landings, San Luis Obispo, (35.17535,-120.72240), intertidal, 2/6/21; TLT1270, Zebra cove, Anacapa Is., (34.01000,-119.44000), 6-13 m, 6/17/22; TLT806, Three sisters, Santa Cruz Is., (33.98675,-119.58884), 6-13 m, 4/16/21; TLT991, Farnsworth Bank, Catalina Is., (33.34380,-118.51650), 21-27 m, 7/10/21; TLT1102, CRABS, Monterey, (36.55377,-121.93840), 10-17 m, 9/21/21; BHAK-03707, Simon’s Landing Drop-down, Tatoosh Island, Washington, (48.391917, - 124.733236), intertidal, 6/15/18; BHAK-03699, Strawberry Draw Overhang, Tatoosh Island, Washington, (48.392003, −124.737667), intertidal, 6/15/18; UF4317, Cape Arago, south Middle Cove, Oregon, (43.30477,−124.40061), intertidal, 6/5/19; UF4300, Cape Arago, Middle Cove, Oregon, (43.30477,−124.40061), undetermined, 6/4/19; BHAK-02365, West Beach, Calvert Island, British Columbia, (51.6495,−128.155), 5 m, 8/2/17; TLT1395, Cape Alava, Washington, (48.17134, −124.75227), 4−12 m, 7/29/23; TLT1386, Cape Alava, Washington, (48.17134, −124.75227), 4−12 m, 7/29/23; TLT1313, Cape Alava, Washington, (48.17134, −124.75227), 4−12 m, 7/29/23; TLT1426, Tatoosh Island, Washington, (48.39370, −124.73305), 4−11 m, 7/31/23.

#### Morphology

Cushion shaped to massive and asymmetrical. Firm and rubbery with velvety surface. Highly contractile; becomes a firm mass without apparent oscula when exposed by the tide. When alive and relaxed, subtidal samples display a partially transparent network of pores on the surface and large, obvious oscula on raised chimneys. Samples vary from yellow to brownish-yellow alive; beige after preservation.

#### Skeleton

Ectosomal skeleton of tylostyles in upright bouquets. Tylostyles in choanosome in confusion with no apparent pattern.

#### Spicules

Tylostyles only. Often straight, but some samples have a high frequency of spicules with slightly curved, bent or sinuous shapes. Heads of California samples are frequently lumpy and uneven (figure 4E), but this was rare in samples from farther North (e.g. British Columbia, Washington). Most samples display a uniform thickness until reaching a sharply tapering point, but some samples slightly fusiform (thickest in center) and/or more gradually tapering. Slightly shorter in the ectosome.

For one sample (TLT296), spicules were measured separately for the ectosome (199–429–559 x 4–7–11 μm (n=124)) and choanosome (261–479–599 x 3–7–12 μm (n=91)) and found to be significantly smaller in the ectosome (Wilcox p-value = 2.6e-07). The combined distribution was not distinctly bimodal due to the similarity of the distributions.

Spicules were measured from 15 California samples, collected from Monterey Bay in Central California to La Jolla cove in Southern California, including 6 different Channel Islands. All samples together: 148–418–615 x 2–7–18 μm (n=570). Mean values varied greatly across samples, with mean lengths from 282 to 511 μm and mean widths from 5 to 12 μm.

Spicules were measured from the holotype and a paratype from British Columbia, and a sample from Tatoosh Island, Washington. Mean lengths of these samples were similar to California samples (387 to 483 μm) but mean widths were slightly larger than the largest California samples (13 to 16 μm).

All measured samples combined: 148–426–615 x 2–9–26 μm (n=748).

#### Distribution and habitat

Known from Queen Charlotte Island in British Columbia to San Diego, CA, with a depth range from the low intertidal to 30 m. Common in the intertidal in northern and central California and probably farther North; southern-most intertidal samples known from Malibu, California. Presence-absence surveys in subtidal kelp-forest habitat in California suggest it is widespread but uncommon; it was seen at 10% of investigated reefs, evenly spread from Monterey to San Diego counties (the entire investigated range). The nudibranch *Doris montereyensis* is frequently observed feeding on this species.

#### Remarks

A species of *Suberites* was first noted in the Central California intertidal nearly 50 years ago, but despite being locally abundant, it has remained unnamed (Hartman 1975; Lee *et al*. 2007). It is fairly common in the intertidal, especially in northern and central California; as such, it is surprising that it was not noted during the extensive intertidal work of de Laubenfels in the 1920s (de Laubenfels 1932). It is possible that de Laubenfels’ subtidal *Ficulina suberea lata* were in fact a mix of *S. latus* and *S. lambei*, but all of those samples were subtidal or in beach wrack rather than intertidal. Consistent with this possibility, de Laubenfels indicated lumpy tylostyle heads in these samples, which were common in our California samples of *S. lambei* (figure 25, de Laubenfels 1932) but not seen in other *Suberites* species we examined.

*Suberites lambei* was described from British Columbia, with the thickness of the spicules as a key trait distinguishing it from *S. latus* (Austin *et al*. 2014). Austin *et al*. compared their new species to a few samples from Central California, but were skeptical that they were the same species because 1) most California samples had much thinner spicules and 2) the populations were disjunct, as they were unaware of samples between British Columbia and Central California. Here, we have collected additional samples from British Columbia, Washington, Oregon, Central California, and Southern California, from both the intertidal and the subtidal.

The spicular dimensions of these samples vary greatly, especially in width, but the continuous distribution of values suggests this is intraspecific variation. DNA data at both *cox1* and *28S* support this assertion (figures 1-2). We therefore have no evidence supporting more than one species among these samples, and assign them all to *S. lambei*.

This species was previously distinguished from *S. latus* by spicule width, but we show that this character is variable. *Suberites lambei* never have microscleres, which differentiated them from all *S. kumeyaay*, all *S. californianai,* and nearly all *S. latus*. As some samples of *S. latus* may lack microscleres, these samples can be morphologically distinuished in the following ways. *Suberites latus* is often orange (*S. lambei* is yellow or light brown); *S. latus* is usually associated with hermit crabs (not seen in *S. lambei*); *S. latus* has a substantial difference in tylostyle length between ectosome and choanosome— nearly twice as long in the choanosome — leading to a strongly bimodal distribution when they are combined (not seen in *S. lambei*). The tylostyles are also usually shorter in *S. latus* than *S. lambei,* but the shortest *S. lambei* sample we investigated was similar to *S. latus* (genotyping confirmed this sample was *S. lambei*), so this character is insufficient in isolation. Finally, we note that these species are differentiated by depth range in California, but not in more Northern parts of their shared range.

***Suberites agaricus* sp. nov. Turner, 2023**

Figures 1-2, 7

**Figure 7.**
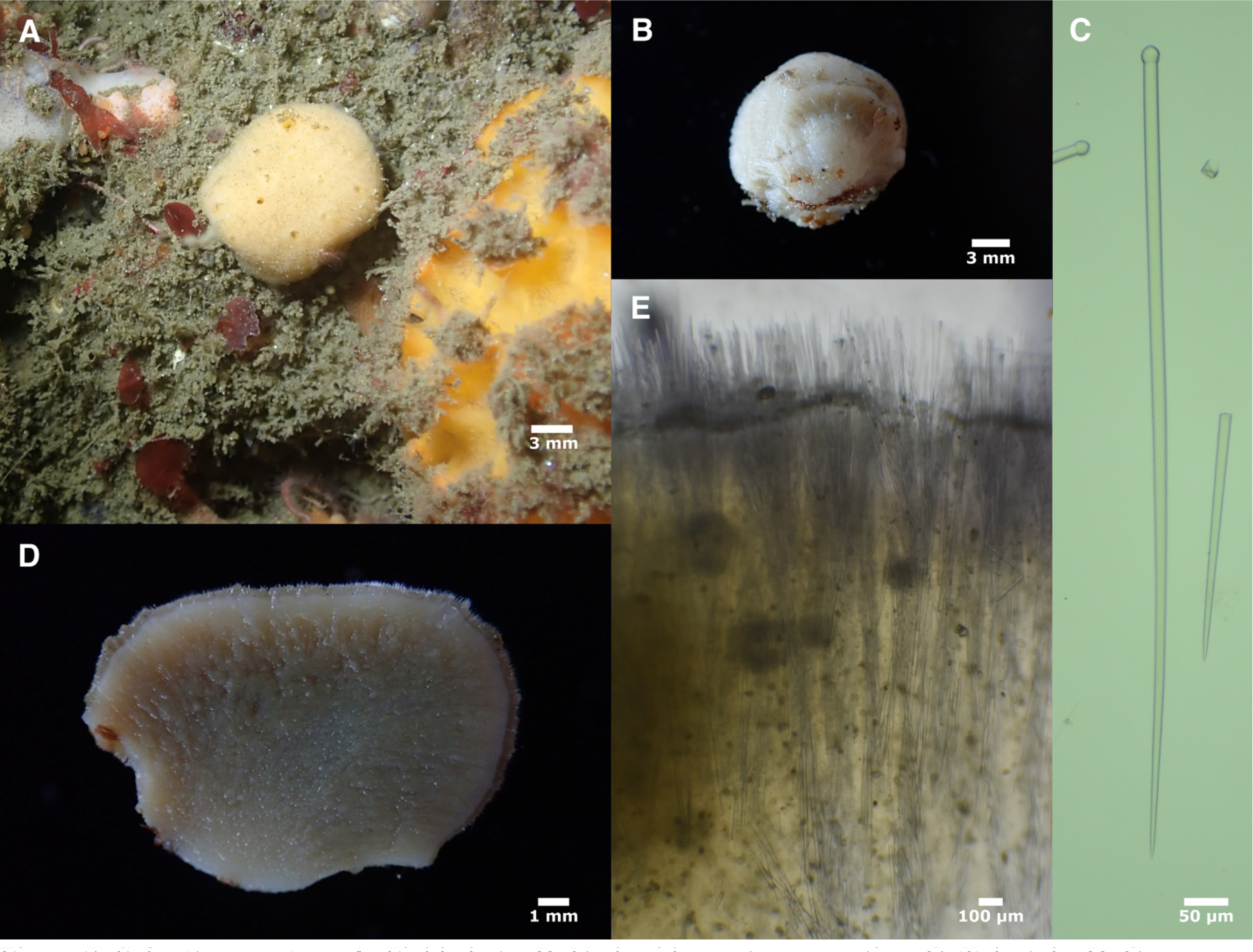
*Suberites agaricus*. A: Field photo. B: Underside, post-preservation. C: Tylostyle. D: Cross-section, post-preservation. E: Skeletal structure in cross-section. All photos of holotype.

#### Material examined

Holotype: TLT1321, El cono, Point Loma, (32.69853, −117.27338), 21-26 m, 5/6/23.

#### Etymology

Named for its resemblance to cultivated “button” mushrooms in the genus *Agaricus*.

#### Morphology

Roughly spherical, 14 mm in diameter. Numerous small oscula (∼1 mm in diameter) flush with surface in living specimen; constricted in preserved specimen. The ventral side appears to have been attached to the substrate over a smaller surface than the entire diameter of the sponge, leading to an incipient stalked appearance, and making the overall morphology similar to the cap of the cultivated “button” mushroom. Pale yellow alive, white when preserved. Cross-section of the preserved sample reveals a visually distinct ectosomal layer 0.2-0.5 mm in thickness (figure 5D). Microscopically hispid.

#### Skeleton

Upright bouquets of tylostyles in the ectosome, with tips protruding 100−150 μm beyond sponge surface. Bouquets are supported by multispicular columns of tylostyles, creating a somewhat radial architecture, becoming increasingly confused towards the interior of the sponge.

#### Spicules

Large and small tylostyles, with overlapping size distributions. The distribution of tylostyle length is bimodal when all tylostyles are considered together, with modes at approximately 350 μm and 1050 μm. If 700 μm is used as a dividing point between classes, then the choanosome contains only long tylostyles, while the ectosome contains 20% long tylostyles and 80% short tylostyles. All tylostyles are straight or slightly curved, usually with symmetrical, round heads, but some heads are sub-terminal. Most spicules are thickest near head and gently tapering throughout their length, but some are slightly fusiform. The gradual tapering differentiates this species from the other *Suberites* found in California, but *R. gadus* is similar.

All spicules together: 204–598–1335 x 6–11–24 μm (n=161), modes at approximately 350 μm and 1050 μm.

Choanosome only: 860–1074–1335 x 11–17–24 μm (n=23)

Ectosome only: 223–548–1073 x 6–9–16 μm (n=87)

Only spicules < 700 μm in length: 204–409–692 x 6–9–16 μm (n=107)

#### Distribution and habitat

Only seen at a single location to date: the kelp forest off of Point Loma, near the southern border of California. Numerous small, roughly spherical sponges matching the morphology of the sampled individual were seen at this location, which were all likely this species.

#### Remarks

This species could arguably be placed in either *Suberites* or *Rhizaxinella*. The body shape and somewhat radial skeleton are more consistent with *Rhizaxinella*, but sponges in this genus are carried on a long, often branching stalks, which are missing in this species. With the added evidence of a close genetic relationship to other *Suberites*, we feel this placement is a better fit. It is likely that the spherical body of the sponge has led to some convergent evolution of the skeleton from confused into a more radial architecture.

This species is quite closely related to *S. lambei*, with only 0.3% sequence divergence at the 28S locus. Divergence at the *cox1* locus is higher (1.7%), and with consistent placement and both loci and very distinct spicule morphology, it is confidently assigned to a separate species. As discussed in the *S. lambei* section, tylostyles in that species have a unimodal length distribution, with average values between 282 μm and 499 μm per sample. We measured 750 spicules across 17 individuals throughout the range of *S. lambei* (including a sample from San Diego, near where *S. agaricus* was collected) and found the maximum spicule length to be 615 μm. This is in stark contrast to *S. agaricus*, where minimum sizes of the long tylostyles are longer than this maximum value for *S. lambei.* These long tylostyles also set the new species apart from all other *Suberites* in the region: *S. latus, S. kumeyaay* sp. nov.*, S. californiana* sp. nov., *S. concinnus* Lambe, 1895 (which lacks tyles), and *S. mineri* de Laubenfels, 1935 (which is cup-shaped). *Suberites baffini* (Brøndsted 1933) has spicules of similar length to *S. agaricus*, and also has a roughly radial architecture. These are unlikely to be conspecific, however, as *S. baffini* is described as having prominent spicular fibers, whereas no spongin or other fiber is present in *S. agaricus*; moreover, *S. baffini* is known from deep water (1200 m) the far North Atlantic (Baffin Bay, Canada).

The spicules of the new species are similar to *Rhizaxinella gadus*. The spicules of *R. gadus* are significantly longer (Wilcox rank-sum test p < 0.001), but examining more individuals may reveal overlap in size distributions. As described in the *R. gadus* section below, these species can be differentiated based on skeletal organization, the lack of a branching stalk in *S. agaricus*, and DNA sequence; they also likely have depth range differences, but further work could reveal overlap in depth range.

**Genus Rhizaxinella Keller, 1880**

***Rhizaxinella gadus* (de Laubenfels, 1926)**

Figures 1-2, 8

**Figure 8.**
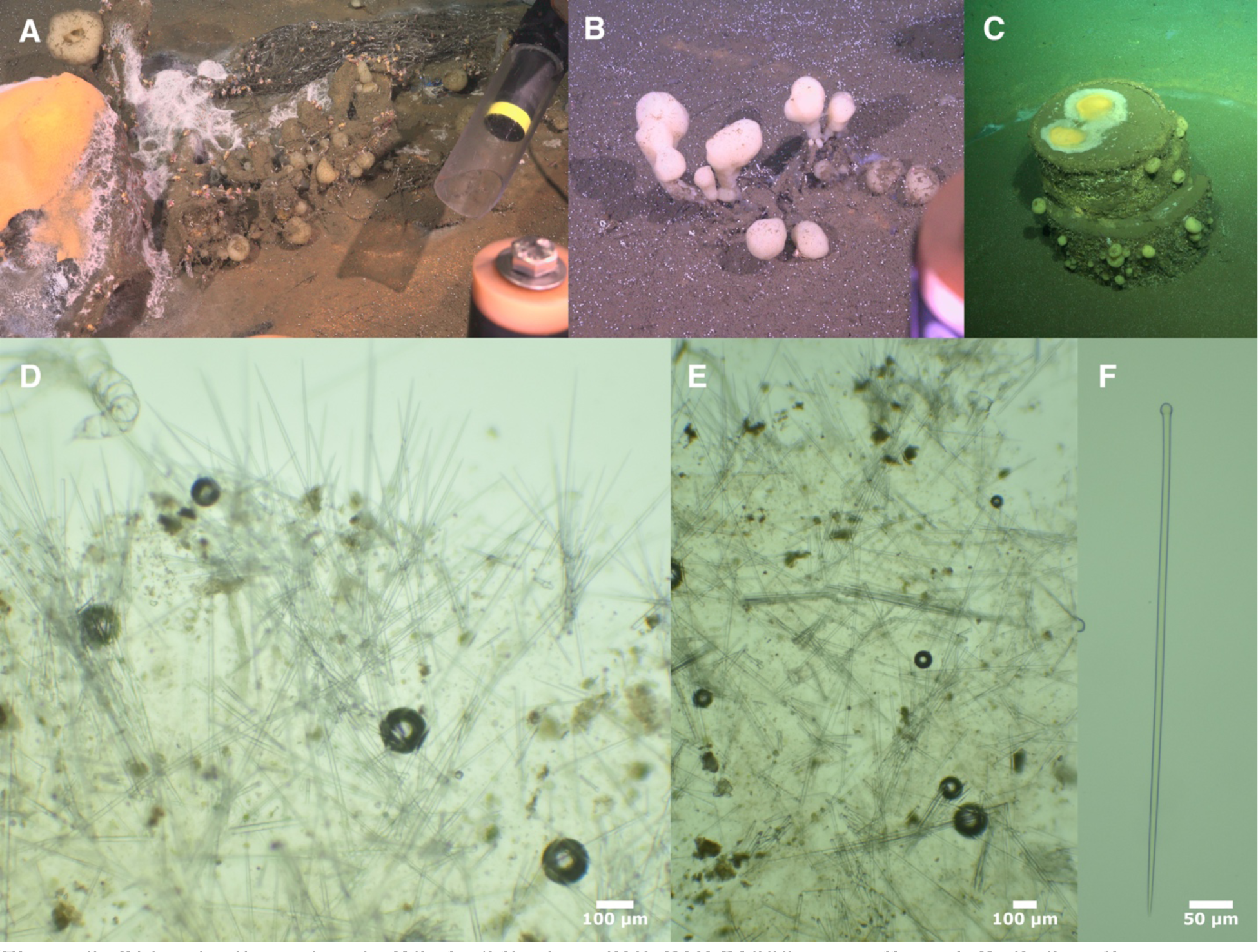
*Rhizaxinella gadus.* A: Whale-fall where SIO-BIC P1999 was collected. B: Soft-sediment where P2000 was collected; when disturbed by the slurp tube, all stalked sponges in this photo were revealed to be connected below the sediment by branched stalks. C: DDT-barrel dump site where SIO-BIC P2015 was collected. D: Cross-section of SIO-BIC P1999 showing ectosomal spicule bouquets and confused choanosomal skeleton. E: Cross-section of SIO-BIC P2016 showing confused spicules and spicule bundles in choanosome. F: tylostyle from SIO-BIC P1999.

### Synonomy

*Suberites gadus* (de Laubenfels 1926, 1932)

#### Material examined

SIO-BIC P1999, San Pedro Basin, whale fall, (33.5736, −118.43676), 885 m, 8/3/21; P2000, San Pedro Basin,, (33.57650, −118.43484), 882 m, 8/3/21; SIO-BIC P2015, barrel 1, San Pedro Basin, DDT barrel dump site, (33.56602, −118.42426), 885 m, 8/4/21; SIO-BIC P2016, barrel 2, San Pedro Basin, DDT barrel dump site, (33.56607, 118.42566), 885 m, 8/4/21. All collected by the Schmidt Ocean Institute using the ROV *SuBastian*.

#### Morphology

Holotype previously described as cylindrical and peanut-shaped sponges atop branching stalks 1 cm thick. Only small fragments of newly analyzed samples were available for morphological analysis, but ROV video from their collection shows globose sponges 2-6 cm in diameter; many were atop thick and sometimes branching stalks, though stalks were not apparent on all sponges. Oscula sometimes visible either as a few scattered holes near the apex, flush with sponge surface, or as a larger single opening, also at apex (oscula not visible in measured samples). White to pale yellow alive; white preserved. Microscopically hispid.

#### Skeleton

Short tylostyles in upright bouquets and bundles in the ectosome, with tips protruding ∼300 μm beyond the sponge surface. Long tylostyles occurring both singly and in bundles of up to 20 spicules in the choanosome. Single spicules and spicule bundles are both found in confusion with no apparent spicule tracts. Samples available for examination did not include stalks; stalks previously described as having a “central axis of densely packed longitudinally placed spicules” (de Laubenfels, 1932) and an ectosomal skeleton the same as the lobes.

#### Spicules

Long and short tylostyles. Most with well-formed, round, terminal heads, but occasionally with subterminal heads or both. Straight or slightly curved; most are gradually tapering throughout their length, but some taper for only half their length or less.

Long tylostyles: nearly all the spicules in the choanosome are in the longer size class, while a minority of spicules in ectosomal spicule preps are also in the larger size class. Distributions are overlapping; if 850 μm is used as the dividing point, then long tylostyles in SIO-BIC P1999 are 852–1076–1512 x 9–12–17 μm (n=74); long tylostyles of SIO-BIC P2016 are 907–1338–1734 x 8–12–17 μm (n=34).

Short tylostyles: Only slightly thinner than long tylostyles, and otherwise with the same shape. Using 850 μm as the dividing point between long and short tylostyles, short tylostyles in SIO-BIC P1999 are 229–521–817 x 5–8–12 μm (n=77); short tylostyles of SIO-BIC P2016 are 260– 462–776 x 4–7–10 μm (n=57).

All tylostyles of SIO-BIC P1999, considered together: 229–793–1512 x 5–10–17 μm (n=151). Distribution is bimodal, with modes at 350 and 1025 μm. Tylostyles of SIO-BIC P2016: 260– 789–1734 x 4–9–17 μm (n=91), with modes at 350 and 1300.

#### Distribution and habitat

The holotype was found near Pacific Grove in Central California; it was tangled in fishing gear and sampled at uncertain depth (depth first reported with skepticism as 180 m and later as 30 m (de Laubenfels 1926, 1932)). Newly collected samples were found on whale falls and on and near metal barrels in the San Pedro Basin of Los Angeles, 880-890 m depth. Some protruded from sediment, with the branching portions of the stalk within the sediment. This species may be common in deep water in the Northeast Pacific, but further samples are needed for confirmation (see remarks).

#### Remarks

This species was described from a single sample found near Pacific Grove, California. Newly collected samples analyzed here match the described sample very well in terms of spicule sizes, spicule morphology, and gross morphology.

There are several hundred records of this species in the Global Biodiversity Information Facility (gbif.org), but nearly all are from the NOAA Deep Sea Corals Research and Technology Program. These records are based on the gross morphology of samples seen on video transects, and were not collected and examined, so they should be considered tentative. We examined the only other sample in the database that was collected in California (CASIZ165849), and found that it did not match the description of *R. gadus* (the sample contained multiple size classes of tylostyles, but they were smaller than *R. gadus*, and the sample also had abundant strongyloxea up to 2500 μm long; only a small fragment was available, so we were unable to determine the identity of this sample, but it is likely from an undescribed deep-water species that may also be in the Suberitidae). It therefore remains possible that there are multiple species of stalked sponges matching the morphology of *R. gadus* in the region, but this remains to be confirmed. The remaining GBIF records are from Mexican waters and were not examined here.

**Genus Pseudosuberites Topsent 1896**

***Pseudosuberites latke* sp. nov. Turner 2023**

Figures 1-2, 9

**Figure 9.**
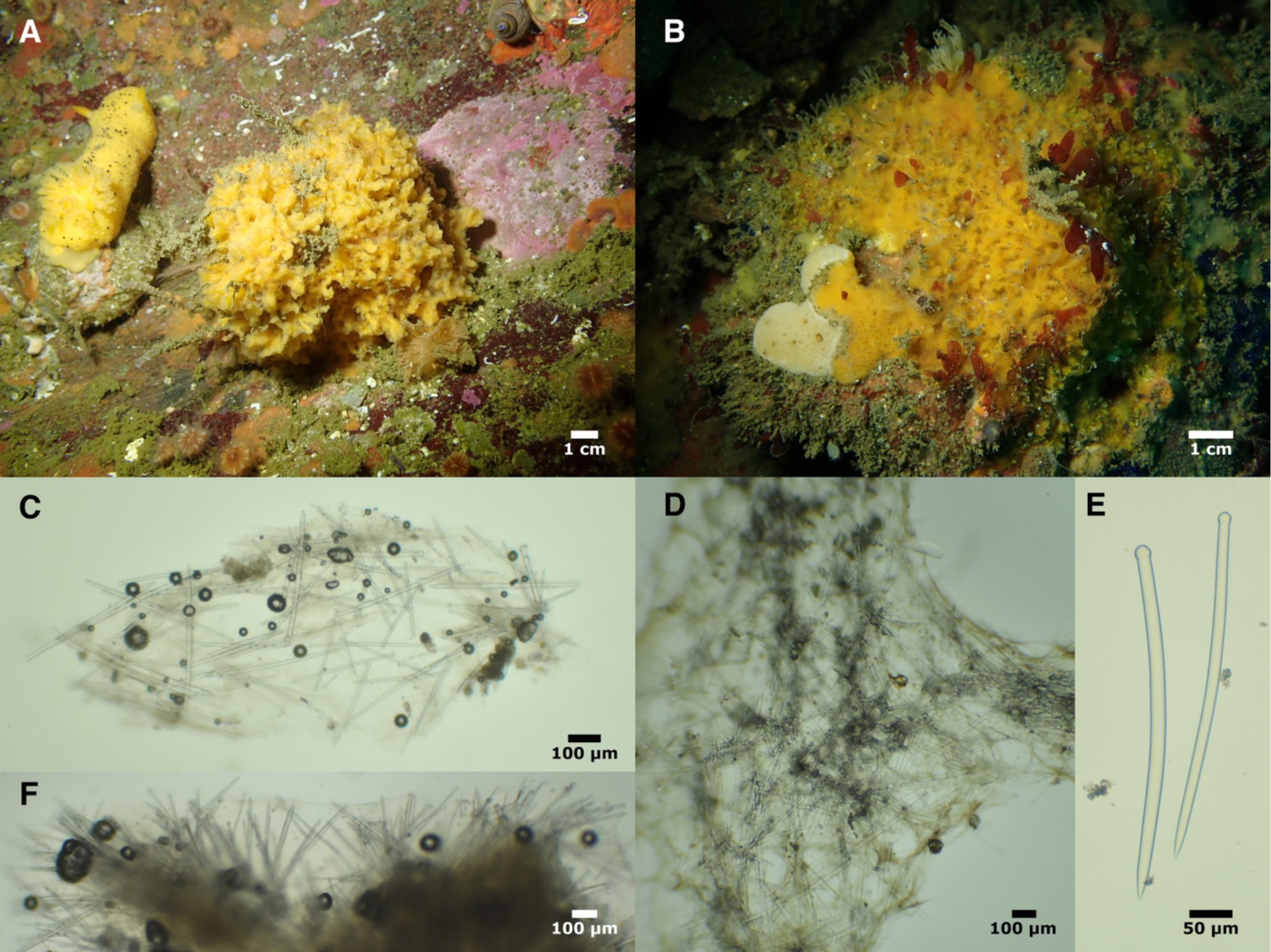
*Pseudosuberites latke*. A: Field photo of holotype. B: Field photo of encrusting sample TLT538. C: Tangential spicules in ectosomal section. D: Dense spiculation of confused reticulation. E: Example tylostyles. F: Partially digested cross-section showing plumose spicules emerging from dense reticulation (which is obscured by remaining tissue).

#### Material examined

Holotype: TLT1131, Fire Rock, Pescadero Point, (36.55898, −121.95110), 10-22 m, 8/10/21. Paratypes: TLT538, Goalpost, Point Loma, (32.69438, −117.26860), 12−15 m, 2/8/20; TLT585, Train wheels, Point Loma, (32.65205, −117.26243), 9/19/20; 15-29 m, 9/19/20; TLT1143A, Tanker Reef, Monterey Bay, (36.60607, −121.88160), 10−16 m, 8/9/21; TLT1128, Elwood Reef, Santa Barbara, (34.41775, −119.90150), 9−15 m, 9/8/21; TLT19, Elwood Reef, Santa Barbara, (34.41775, −119.90150), 9−15 m, 4/17/19; TLT433, Elwood Reef, Santa Barbara, (34.41775, −119.90150), 9−15 m, 10/23/19.

#### Morphology

Holotype is a bushy mass, 30 mm thick 8 mm across, composed of a reticulate network of individual branches 2-5 mm in diameter. Other samples similar, but some encrusting instead of bushy, 5 mm thick, with the reticulate mass prone on substrate rather than upright. Bright yellow alive, white preserved. Oscula sometimes visible in photos of living sponges.

#### Skeleton

A dense confused mass of tylostyles is formed into a reticulate network. Plumose bundles and individual spicules protrude from this confused mass and pierce the surface of the sponge, making ectosome difficult to detach. Ectosomal sections contain tangential tylostyles in confusion.

#### Spicules

Tylostyles only. Highly variable in length and width, but without bimodal distributions. Nearly all with well-formed, round, terminal heads, but occasionally with subterminal heads. Straight or slightly curved, with uniform thickness along shaft until reaching a sharply tapering point.

Holotype: 247–400–622 x 6–12–23 μm (n=131). When ectosome and choanosome are prepared separately, spicules in ectosome are significantly shorter (p=0.016), but distributions are broadly overlapping (ectosome only: 247–361–473 x 6–10–15 μm (n=31); choanosome only: 263–410– 622 x 6–13–23 μm (n=42)).

All samples combined: 176–388–730 x 2–11–23 μm (n=265). Means of each sample vary from 341 to 420 μm in length, 6 to 12 μm in width.

#### Distribution and habitat

All known samples were recently collected by one of the authors. Presence-absence surveys in subtidal kelp-forest habitat in California found this species to be uncommon from Monterey to San Diego (the entire investigated range); the shallowest samples were found at 10 m and the deepest at 30 m (the maximum depth investigated). It was most abundant in southern San Diego County, where it was seen at 5/12 sites investigated. Many of these were deeper (20-30 m) sites, where it may be more abundant than in very shallow water. Not seen at any island sites, intertidal sites, or on human structures. The nudibranch *Doris montereyensis* has been observed feeding on this species.

#### Remarks

*Suberites* and *Pseudosuberites* are currently distinguished by their ectosomal skeleton (a tangential and usually detachable crust carried by subectosomal spicule brushes in *Pseudosuberites*, a palisade of small tylostyles in *Suberites*) and their tylostyle size distributions (a smaller ectosomal size class in *Suberites* and no localized size classes in *Pseudosuberites*). This species did have some tangential ectosomal spicules, though the surface was not easily detachable as it is in some *Pseudosuberites*. Two size classes vs. one size class of tylostyles is likely not a discrete character, as species like *P. latke* sp. nov. and *S. lambei* have no size classes but significantly smaller spicules in the ectosome. Combining DNA and morphological data leads us to assign this species to *Pseudosuberites*. It is very distantly related to all genotyped *Suberites*, but forms a clade with *Pseudosuberites* from South Africa, Antarctica, and Korea at the *cox1* locus.

The dense reticulation of confused tylostyles makes this species very distinct from any other species known from the region. No other *Pseudosuberites* are described from the Northeast Pacific. The type species *P. hyalinus* was once said to have been collected in Pacific Mexico (Dickinson 1945); little description is provided, but the spicules are listed as three times as long as those of *P. latke* sp. nov.. A more recent publication on the sponges of the tropical Mexican Pacific includes a “*Pseudosubertes* sp.*”* but no morphological description is yet available (Carballo *et al*. 2019).

**Genus Protosuberites** Swartschewsky, 1905

***Protosuberites sisyrnus* (de Laubenfels 1930)**

Figure 10

**Figure 10.**
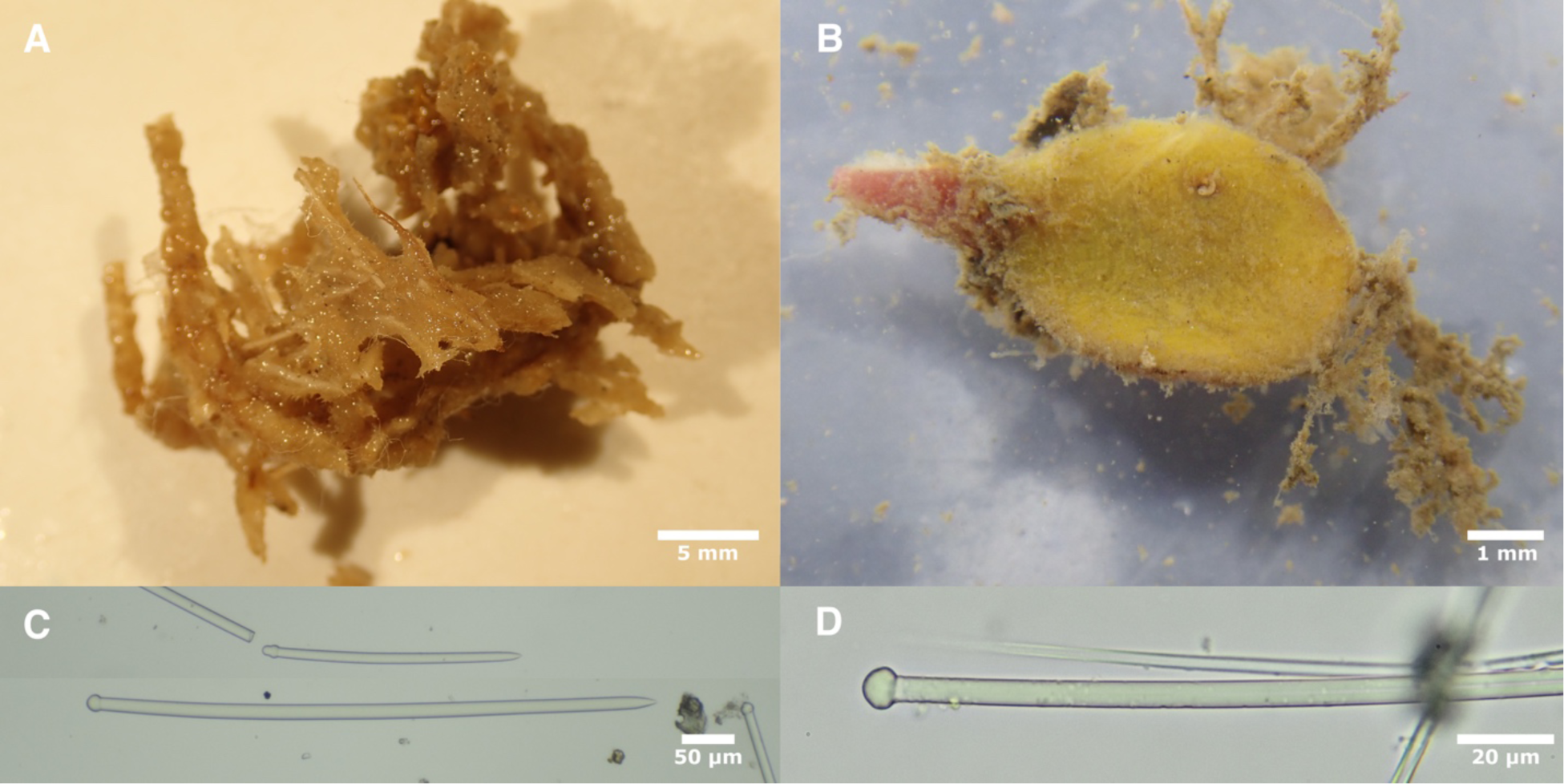
*Protosuberites.* A: *Protosuberites sisyrnus* holotype fragment post-preservation. B: *Protosuberites sp.* TLT443 before preservation on day of collection. C: Long and short tylostyles from *P. sisyrnus*. D: Example tylostyle head from *Protosuberites sp.* TLT443.

#### Material examined

Fragment of holotype, found in the Natural History Museum Los Angeles collection, labeled as “21413 (Part of type)”, with collection information that fits holotype. The holotype is also stored in the Smithsonian as USNM21413. Collected South of the breakwater, San Pedro, California, 45 m, 4/5/1924.

#### Morphology

Sponge is encrusting on a tangled mass of polychaete tubes and other debris. Visibly hispid. Difficult to further characterize due to the age and complexity of the sample. Light brown post-preservation.

#### Skeleton

Not investigated; previously described as erect tylostyles in the ectosome and few spicules, in confusion, in the choanosome.

#### Spicules

Tylostyles; both straight and slightly curved spicules are common, most with round terminal heads, but occasionally subterminal. Consistent thickness through shaft until reaching a sharply tapering point. Considerable variation in length and width but distribution unimodal. 246–386–574 x 7–12–19 μm (n=50)

#### Distribution and habitat

Described from two samples dredged from 45-54 m near Los Angeles and Catalina Island in Southern California. Museum records indicate occurrences in British Columbia, but this remains to be confirmed, as samples we examined were misidentified (see remarks).

#### Remarks

We were unable to locate any samples of this species in the field (previously collected samples were from a depth range we were not able to investigate). Vouchers previously identified as *Protosuberites* were loaned to us from British Columbia, but these proved to be misidentified (they are likely in the Microcionidae, as spicules consist of palmate chelae, toxas, and two sizes of acanthostyles). We attempted to sequence DNA from several loci from the holotype fragment in the Natural History Museum of Los Angeles but were unsuccessful. The quantitative spicule data presented here should aid in future attempts to locate and characterize this species.

***Protosuberites sp.***

Figure 1-2, 10

#### Material examined

TLT443, Elwood Reef, Santa Barbara, (34.41775, −119.90150), 9−15 m, 10/23/19.

#### Morphology

Thinly encrusting on a small (∼1 cm) blade of red algae. Bright yellow alive.

#### Skeleton

Not investigated; the entire sample was consumed for DNA extraction and spicule preparation.

#### Spicules

Tylostyles, straight or weakly curved. Often with well-formed terminal heads, but a high frequency of heads that are sub-terminal or slightly lumpy. Longer spicules are gently tapering for much of their length, but shorter spicules have a consistent width until tapering sharply at tip. Highly variable in length, but distribution is unimodal. 103–246–474 x 2–4–7 μm (n=43).

#### Distribution and habitat

Found at a single location, Elwood Reef, a shallow, rocky reef with kelp forest cover near Santa Barbara, California.

#### Remarks

Thinly encrusting Suberitidae are currently placed in either *Terpios* or *Protosuberites* (Morrow & Cárdenas 2015; van Soest 2002). Though we did not examine the skeleton of this small sponge, we think it is more likely to be in *Protosuberites* due to the shape of the tylostyles and the lack of gelatinous consistency. DNA supports this placement, as the species falls into a clade containing a worldwide sample of other *Protosuberites*.

With only a single sample that was consumed without investigating the skeleton, and noting how variable we found spicule lengths to be in *S. lambei*, we conservatively leave this sample as an unknown for the time being.

**Genus Homaxinella**

***Homaxinella amphispicula* (de Laubenfels 1961)**

Figures 1-2, 11

**Figure 11.**
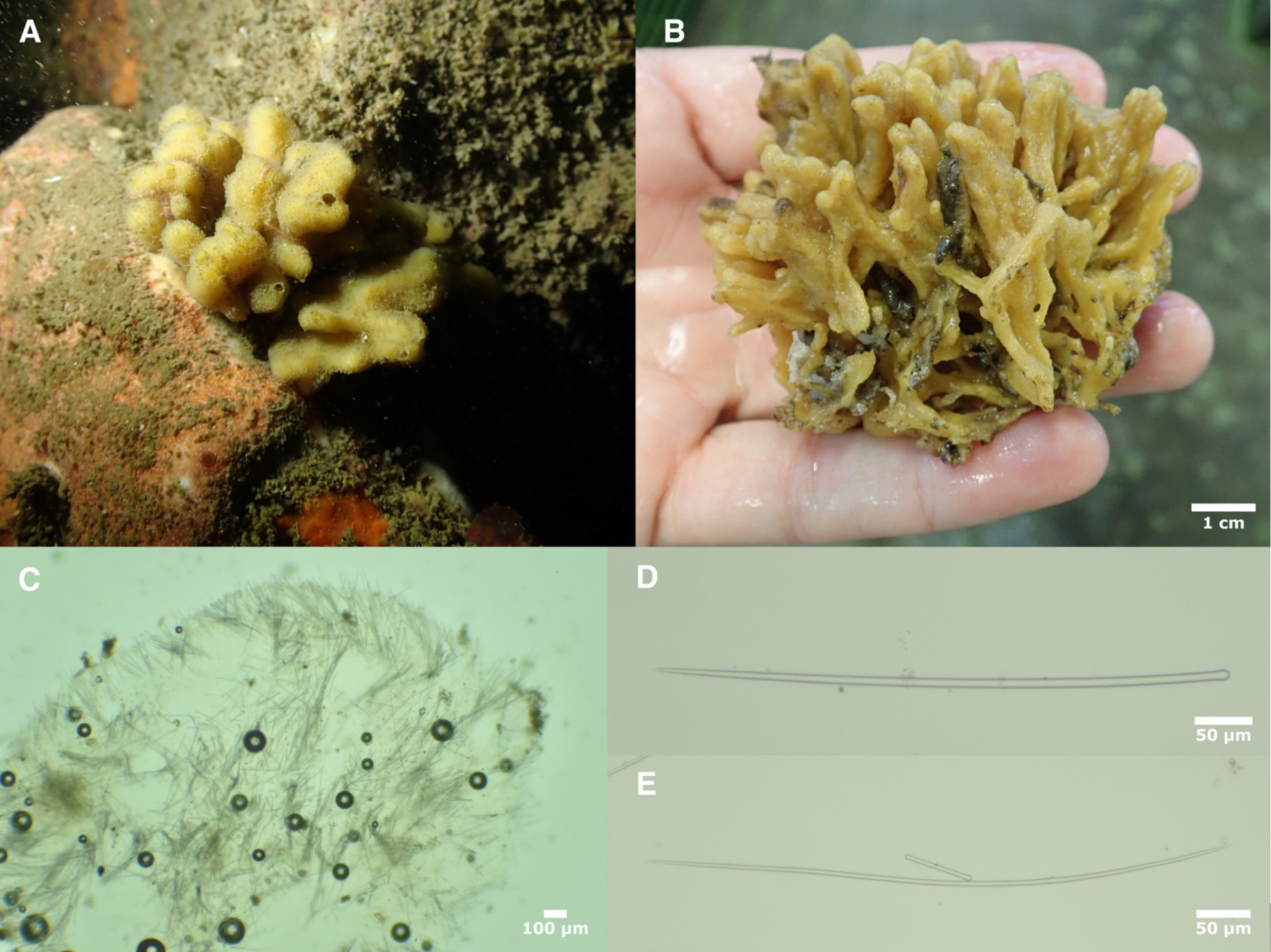
*Homaxinella amphispicula.* A: Field photo of TLT1137. B: Sample TLT1177 before preservation on day of collection. C: Cross section of TLT1177. D: Style from TLT1177. E: Oxea from TLT1177.

### Synonomy

*Syringella amphispicula* (de Laubenfels 1961)

#### Material examined

TLT1137, Elwood Reef, Santa Barbara, (34.41775, −119.90150), 9−15 m, 9/8/21; TLT1177, Elwood Reef, Santa Barbara, (34.41775, −119.90150), 9−15 m, 9/1/21; RBC018-00132-001, Sechelt Peninsula; Sakinaw Lake, British Columbia, (49.56760, - 123.80415), 0−15 m, 7/23/15.

#### Morphology

Upright, branches 2−10 mm in diameter split and anastomose to form a bush mass. Occasional oscula visible at apex of branches in living specimen; contracted and invisible after collection. Dull brownish-yellow alive, beige preserved.

#### Skeleton

Spicule bundles run parallel to branches in the core of each branch. additional bundles, and single spicules in confusion, radiate out from core to periphery. Upright spicules pierce the surface at the periphery in disorganized bouquets and individually.

#### Spicules

Long thin styles, slightly curved or rarely strongly curved or sinuous; heads vary from simple styles to subtylostyles, with intermediates. Highly variable in length, but not falling into easily delineated size classes. Significantly smaller in ectosome, with longest styles (>515 μm) not found in ectosome. Both California samples also had oxeas, but they were rare (<1% of spicules).

Styles of California samples: 197–422–737 x 2–6–12 μm (n=171); ectosome only of sample TLT1177: 224–338–513 x 2–4–7 μm (n=44).

British Columbia sample with slightly longer and much wider styles: 288–525–733 x 6–12–17 μm (n=38).

Oxeas of California samples: 357–536–687 x 3–5–7 μm (n=12).

#### Distribution and habitat

Previously known from the San Juan Islands in Northern Washington to British Columbia, 10-70 m in depth (Austin *et al*. 2014). New samples from Santa Barbara California are a Southern range extension of over 1,500 km.

#### Remarks

As this species was previously known only from British Columbia and the San Juan Islands, we were surprised to find it in Santa Barbara. The first sample was collected by Christoph Pierre and Christian Orsini (UCSB), and a second sample was collected soon after, at the same location, by one of us (T. Turner). Both California samples have rare oxeas, which were not found in the large number of British Columbia samples previously analyzed (Austin *et al*. 2014), nor were we able to find them in the British Columbia sample we analyzed. California samples also have much thinner spicules than British Columbia samples (but we note that thickness is variable in other species, and is sometimes associated with water temperature or chemistry). We were able to generate DNA data at the 28S locus for one British Columbia sample and one Santa Barbara sample, which differ by 0.3%; these samples differ from the North Atlantic *H. subdola* (Bowerbank 1866) by 0.8%. These limited DNA data are therefore consistent with both conspecificity or closely related taxa. As the morphological differences are slight, we conservatively assign the California samples to *H. amphispicula*, but note that further work may reveal them to be a different species.

### Dichotomous Key to California Suberitidae

1A. Macroscleres are tylosyles; may also have centrotylote microstrongyles …2

1B. Macroscleres are styles and/or subtylostyles, with or without oxeas: *Homaxinella amphispicula* (de Laubenfels 1961)

2A. Thickly encrusting to massive (more than 2 mm thick) …3

2B. Thinly encrusting (less than 2 mm thick): *Protosuberites* (*P. sisyrnus* (de Laubenfels 1930) is the only described species, but a second species of *Protosuberites* may be present)<colcnt=3>

3A. Some tylostyles over 800 μm in length …4

3B. All tylostyles less than 800 μm in length …5

4A. Globular sponges atop branching stalks; choanosomal skeleton consisting of spicules and spicule bundles in confusion; collected in deep water: *Rhizaxinella gadus* (de Laubenfels 1926)

4B. Sponge unstalked; choanosomal skeleton containing multispicular columns; collected in shallow water: *Suberites agaricus* sp. nov. Turner 2023

5A. Choanosomal skeleton a dense, confused mass of tylostyles formed into a reticulate network; ectosome with only sparse brushes of upright tylostyles and some tangential tylostyles: *Pseudosuberites latke* sp. nov. Turner 2023

5B. Choanosomal skeleton confused and/or containing radial tracts; ectosome packed with numerous upright tylostyles in a palisade or in bouquets …6

6A. Containing centrotylote microstrongyles: …7<colcnt=3>

6B. No centrotylote microstrongyles …9

7A. Maximum tylostyle length < 500 μm…8

7B. Maximum tylostyle length > 500 μm; sponge sampled from deep water; associated with hermit crab; centrotylote microstrongyles densely packed in ectosome & less common in chaonosome: *Suberites californiana* sp. nov. Turner 2023

8A. Centrotylote microstrongyles densely packed in ectosome & less common in chaonosome; sponge sampled from shallow water; not associated with hermit crab: *Suberites kumeyaay* sp. nov. Turner 2023

8B. Centrotylote microstrongyles more common in choanosome or very rare in both choanosome and ectosome; sponge sampled from deep water; associated with hermit crab: *Suberites latus* Lambe 1893 (in part)

9A. Tylostyles average over 350 μm in length and/or 10 μm in width: *Suberites lambei* Austin, Ott, Reiswig, Romagosa & McDaniel, 2014 (in part)

9B. Tylostyles average less than 350 μm in length and less than 10 μm in width …10

10A. Sponge sampled from deep water; associated with hermit crab; maximum tylostyle length < 500 μm; tylostyles with well-formed heads: *Suberites latus* Lambe 1893 (in part)

10B. Sponge sampled from the shallow subtidal or intertidal; not associated with hermit crab: *Suberites lambei* Austin, Ott, Reiswig, Romagosa & McDaniel, 2014 (in part)

## Conclusions

Here we present the most comprehensive investigation of the Suberitidae of California to date. We have resolved long-standing mysteries (the identity of the intertidal *Suberites* sp.) and discovered and described several new species. By combining DNA and morphological analyses on the same samples, we show that species within the family can be identified morphologically within a regional fauna. This finding is consistent with recent revisions in other regions that used morphological characters alone and were able to discover and describe new species in the family (Cóndor-Luján *et al*. 2023; Fortunato *et al*. 2020; Morozov *et al*. 2018). However, our molecular phylogeny indicates that most of the genera within the family are polyphyletic. For example, a very deep divergence within the genus *Suberites* indicates that species of *Suberites* associated with hermit crabs share a more recent common ancestor with *Halichondria panicea* than they do with species like *Suberites lambei*, *S. diversicolor*, and *S. aurantiacus*. Resolving these challenges will likely require DNA data from type species from additional genera, such as *Rhizaxinella pyrifera* (Delle Chiaje 1828). Once these data are available, however, it seems likely that traditional taxonomic characters will be insufficient to create a taxonomy consistent with evolutionary history. In species with evolutionarily conserved yet ecologically plastic morphologies, it seems likely that biochemical traits will be needed, and we propose that DNA-based characters are likely to be the most easily quantified biochemical traits available (e.g. Lawley *et al*. 2021). It is therefore preferable that future taxonomic work in this family include DNA sequencing, and, at a minimum, crucial that samples be collected and preserved in ways that allow future investigators to extract and sequence DNA.

## Acknowledgements

We are grateful for the help and support of many people in UCSB’s Marine Science Institute and Diving & Boating Program, especially Robert Miller, Clint Nelson, Christoph Pierre, and Christian Orsini. NOAA provided the R/V Tegula small boat support for diving operations, and Steve Lonhart (NOAA) and Shannon Myers (UCSC) were instrumental in facilitating collections in Central California. The Natural History Museum of Los Angeles’ DISCO program facilitated collections in Los Angeles County. Hugh MacIntosh, Christina Piotrowski, Kathy Omura, and Charlotte Seid graciously provided access to their collections at the Royal BC Museum, California Academy of Sciences, the Natural History Museum of Los Angeles, and the Scripps Institution of Oceanography, respectively; Megan Lily, and Wendy Enright (City of San Diego) and Brandon Stidum also provided crucial samples. Several of the California Academy of Sciences samples were collected by the Coral Reef Research Foundation under contract to the US National Cancer Institute. Thanks also to the Schmidt Ocean Institute and chief scientist Lisa Levin, and the science party of the R/V *Falkor* cruise FK210726. Many thanks to the pilots of the remote operated vehicle *SuBastian* for crucial assistance in collecting *Rhizaxinella gadus*. We also thank Matt Whalen, Margot Hessing-Lewis, and the staff, students and volunteers of the Hakai Institute for support during the 2017 Hakai-MarineGeo bioblitz, and the Makah Tribal Nation for supporting the study of sponges from Tatoosh Island.

